# A novel mathematical construction for identifying attractors from task-driven fMRI data

**DOI:** 10.64898/2026.02.13.705859

**Authors:** Hassan H. Abdallah, John Kopchick, Joseph Hadous, Philip Easter, David R. Rosenberg, Jeffrey A. Stanley, Andrew Salch, Vaibhav A. Diwadkar

**Author notes:** Correspondence Dept. of Mathematics, Applied Physics & Mathematics, RM 7101 University of California, San Diego, La Jolla, CA 92093.

## Abstract

Functional brain imaging data can provide a window into the task-driven network states that shape brain function (or dysfunction). Conventionally, these network states can be represented as bivariate correlation matrices (which are formed from fMRI time series from multiple brain regions/nodes within any task window). Here, we treat these conventional connectivity matrices as *connectivity terrains* in order to recover local structure. In principle, any such terrain can be traversed node by node, where from any node, one can move towards its nearest functional neighbor (i.e., its maximally correlated node). In terrains with meaningful structure, such traversals across multiple nodes should converge to *attractor nodes;* here, the nodes that flow into a shared attractor form an *attractor basin,* which effectively is a sub-network within the system.

Extant methods (e.g., degree distribution and characteristic path length) can summarize global network properties but cannot identify attractor nodes and basins. Here, we construct a new relation, called *transitive maximal correlation (TMC)* that can recover attractors and attractor basins in connectivity terrains. Node A is said to be transitively maximally correlated to node B *if and only if* B is an attractor *into which* A flows. We first develop the mathematical basis for deriving a TMC matrix TMC(M) from a bivariate correlation matrix M (before explaining this with hypothetical data). We next apply the TMC relation to connectivity terrains derived from real fMRI time series data, where these data were acquired in two distinct task-domains (that varied in their extent of cross-cerebral demand): i) associative learning and ii) visually guided motor control. We show that TMC is remarkably sensitive to inter-hemispheric structure in the connectivity terrain; here, attractor pairs that were inter-hemispheric homologues were more likely to be observed for the cross-cerebral learning task, than the more circumscribed motor-control data. We confirm the condition-specific sensitivity of TMC showing that observed attractor basins differed significantly across conditions of the learning task. Finally, we demonstrate that TMC complements graph theoretic constructions like path length and betweenness centrality. We suggest that TMC is a mathematically sound and novel method for capturing functional properties of brain networks.

## 1. Introduction

Bivariate correlation matrices provide vivid information about functional relationships between the components of any large-scale complex system [1]. In functional neuroimaging data, these matrices are typically formed after the statistical analyses of the similarities between pairs of time series (usually derived from clearly delineated brain regions)[2–4]. Bivariate correlation matrices are highly informative precisely because the system’s functional states are embedded within them. This embedded functional information can be extracted using a variety of methods including high dimensional machine-learning based classification [5–8], graph-theoretic summarization to capture global and regional connectomic information [9, 10] and inferential statistics (to identify significantly “connected” or “disconnected” pairs of regions)[11, 12].

### 1.1. Connectivity terrains

In the simplest of terms, bivariate correlation matrices can be construed as connectivity terrains, where the terrain has both local and global structure and connectivity gradients can be used for functional parcellation [13]. Thus, each brain region can be thought of as a point in the terrain, and each node’s functional relationships with all other vertices endow the region with *local* functional structure. In principle, this terrain can be traversed one node at a time; in any space with *m* nodes (*N*_1_-*N_m_*), we can begin with *N*_1_ and then move towards *N*_1_’s nearest functional neighbor (i.e., the node with which it is maximally correlated). This process can be repeated till we arrive at a node from where no further movement is possible. If we imagine this terrain in such a way that the direction of steepest descent is always the direction of the most highly correlated node, then when the path ends, the node at the terminus of the path lies at a local minimum of elevation (i.e., it lies at a trough in the functional terrain). This terminal node can be conceptualized as an “attractor” for *N*_1_, because within the terrain, *N*_1_’s functional path converged to it.

In a bivariate correlation matrix, we will show that rather than ending on a single node, this traversal instead ends at a pair of nodes (“attractor pairs”) that are themselves mutually *maximally correlated*. In other words, our formulation defines attractors not as single nodes but as pairs of nodes.

### 1.2. An illustrative example

Consider the bivariate correlation matrix (Figure 1a) derived using time-series from 15 brain regions (r_1_ - r_15_). Following the discussion above, Figure 1b schematically renders this matrix as a functional terrain. In this terrain, edges are drawn between nodes and their nearest functional neighbor. Nodes are arranged such that each edge traverses a path of steepest descent. As seen, the fifteen regions in this terrain eventually flow into two attractor pairs (best visualized in a side view). Thus, based on this common property, the fifteen regions can be seen to form two functional neighborhoods, which we term *attractor basins*. These basins are “local” *sub-networks* that can be identified using a simple procedure applied to bivariate correlation matrices.

**Figure 1.**
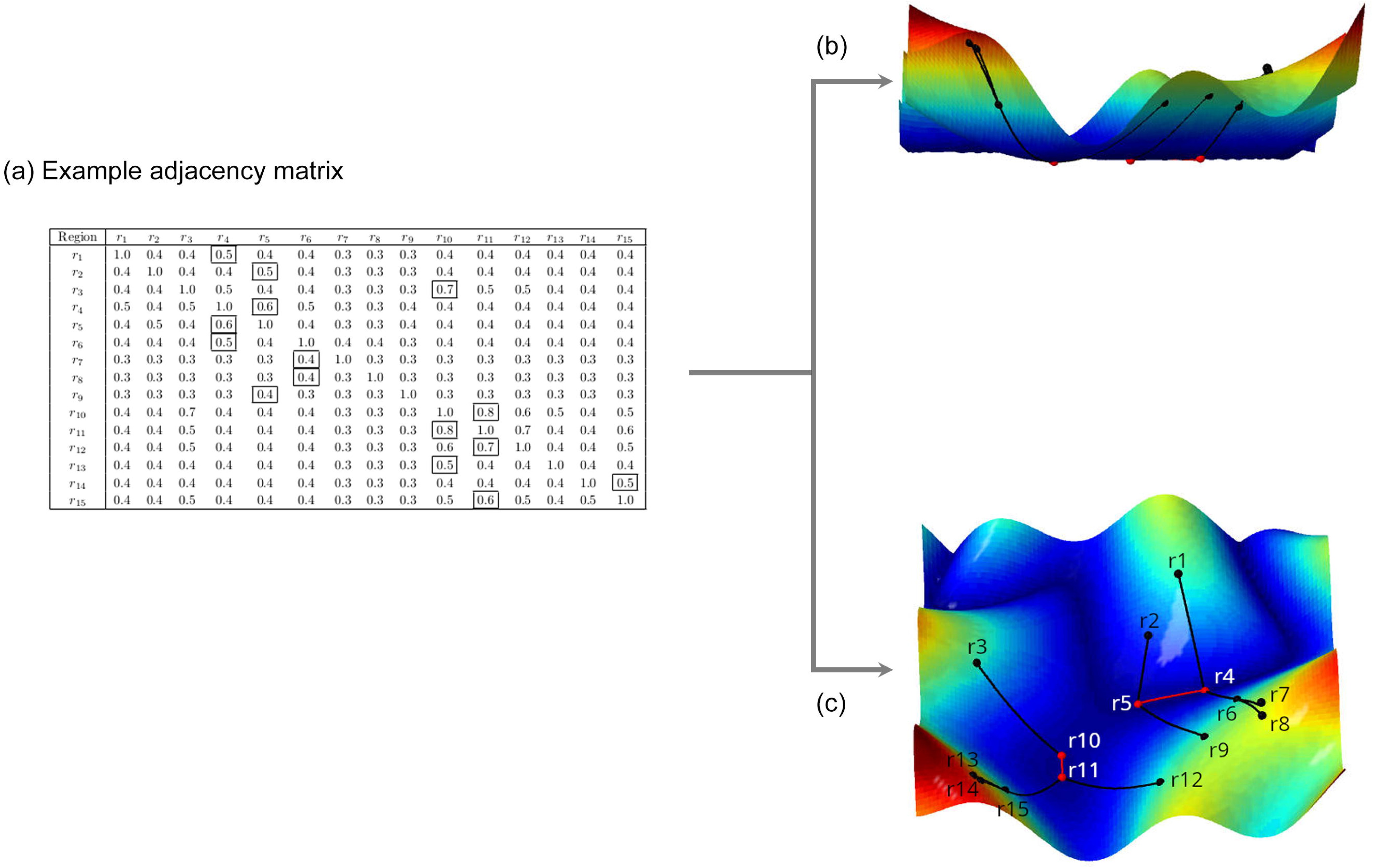
This example illustrates a recurring theme used in this paper: That adjacency matrices (a) can be visualized as connectivity terrains, two views of which are depicted in (b) and (c). These connectivity terrains can be traversed. As we move from a given region to another region that it is transitively maximally correlated to, we envision that we are moving downhill on a surface along a path of steepest descent leading to attractors in the terrain. (a) An example bivariate correlation matrix is provided for a fifteen-region space. The largest non-diagonal entry in each row is marked with a box, because these entries carry the data denoting the regions which are connected by flow lines in (b) and (c). For example, since the highest non-diagonal entry in row 1 is in column 4, region 1 flows into region 4. (b) and (c) are two views of the connectivity terrain derived from the adjacency matrix. The red and black dots are intended to depict regions, where the red dots are attractors. A black path is a “flow line,” and is drawn from a region *r*_n_ to another region *r*_m_ if and only if *r*_m_ has the highest correlation with *r*_n_ among all regions other than *r*_n_ itself. Thus, we see that this correlation matrix yields two attractor basins, and each basin flows into an attractor *pair*, i.e., a pair of regions which mutually flow into one another. An interactive version of the terrain is available at https://hassanabdallah.com/plots/connectivity_terrain.html.

Below, we provide a novel mathematical construction for identifying attractors and attractor basins from task-driven fMRI data. Working with connectivity terrains based on bivariate correlation matrices, we develop the notion of *transitive maximal correlation* (TMC): thus, node N_i_ is transitively maximally correlated to node N_j_ if and only if N_j_ is an attractor into which N_i_ “flows”. An *attractor basin* is the set of all nodes that are transitively maximally correlated to a given attractor pair. We position this method as a complement to methods that are based on global summaries (e.g., graph theoretic methods), such as degree distribution and/or characteristic path length, or betweenness centrality, and provide a formal approach below. These global methods are designed to quantify the integrative roles of nodes *across* the entire set of network edges [14], rather than being suited for identifying local structures such as attractor nodes or basins [15, 16].

### 1.3. A formal approach

Suppose we are given a bivariate correlation matrix *M* that summarizes functional similarities between all *N* constituents (regions or nodes) of a system (brain). *M* represents pairwise functional similarities across all possible pairs of *N* brain regions (i.e., _N_C_2_ unique pairwise relationships).

Now, write *S* to denote the constituents of the set from which the elements of the matrix *M* are composed. Thus, *S* is a set of *N* random variables, and we would like to speak of *sub-networks* within this set *S* of random variables. The question is: what is the simplest procedure to identify sub-networks inside of *S,* from the contents of the correlation matrix *M*?

We address this problem in an axiomatic way. Here are two reasonable axioms which we assume, in order to make any talk of sub-networks in *S* conform to some basic intuitions about what a “network” ought to be:

1. A sub-network in *S* is a subset of *S*.
2. Suppose that *x,y* are members of *S,* such that *y* is maximally correlated to *x*.

Then every sub-network in *S* containing *x* also contains *y*.

For concreteness, we spell out what these axioms mean in the context of summarized fMRI data:

1. A sub-network is a set of regions of interest.
2. If *r_1_, r_2_* are regions of interest, and *r_2_* is more strongly correlated with *r_1_* than with any other region besides *r_2_*, then every network containing *r_1_* also contains *r_2_*.

Clearly these axioms do not, on their own, uniquely determine the sub-networks in *S*. For example, we could simply say that *S* itself comprises a single network and cannot be further parcellated into additional sub-networks. This satisfies the two above axioms. Hence, we can satisfy the above axioms in a trivial way by declaring our sub-networks to be *as large as S* itself.

Consider the opposite case: what sub-network partitions are observed if we ask for each sub-network in *S* to be as *small* as possible, while still satisfying the two axioms? We answer this with the following theorem (whose proof is presented in Appendix 4):

**Theorem**. *Given a member r of S, there is a unique smallest sub-network in S containing r. This smallest sub-network is precisely the attractor basin containing r*.

Even though this theorem is quite easy to prove, its consequences are important: the attractor basins, which we defined entirely in terms of transitive maximal correlation, and without any reference to any particular notion or theory of “networks,” in fact yield a *canonical* theory of networks associated to any bivariate correlation matrix. If we adopt the two axioms above, *then the smallest possible sub-networks among the random variables are precisely the attractor basins*.

In what follows, we further develop the mathematical basis for deriving a TMC matrix *TMC(M)* from a bivariate correlation matrix *M*. Next, we apply the construction to bivariate correlation matrices derived from fMRI time series data acquired in two distinct task-domains: a) associative learning, a task which evokes large-scale interactions between regions in the sensory cortex, the medial temporal lobe and regions in the hetero-modal cortex [11, 17–20] and b) visually guided motor control, a task that evokes more circumscribed interactions between regions in the visual, pre-motor and motor cortices [21, 22]. From these applications, we demonstrate that our method is sensitive in identifying task-specific attractor characteristics. For example, we will show that attractor pairs that are identified from the learning data are more likely to be inter-hemispheric homologues, than attractor pairs that are identified from the motor control data. Results like these reveal a rich set of properties identified by TMC, including the task- and condition-related specificity of attractors, and the observation that transitive maximal correlation *is not predicted by the anatomical proximity* of brain regions. In subsequent discussion, we suggest that TMC is a novel and reliable construct for capturing local functional properties of neuroimaging data from bivariate correlation matrices and should be used to discover functional properties of data across tasks and study populations.

## 2. Methods

### 2.1. The transitive maximal correlation matrix *TMC (M)*

In this section we describe the mathematical process for deriving a *transitive maximal correlation matrix* from a bivariate correlation matrix. While our motivating application is for task-based fMRI, the application could be extended to any class of data representing mass bivariate relationships between pairs of constituents in any dynamical system.

Suppose we are given a symmetric square matrix *M* of real numbers. In the area of intended application of these ideas, we would fix an experimental condition γ, and we would have some set {*x*_1_,*x*_2_,…,*x_n_* } of regions of interest in the brain. The entry in row *r* and column 𝑐 of the matrix *M* should measure the strength of the relationship between the fMRI time series data from region *x*_*r*_ and region *x*_𝑐_ during condition γ (i.e., the absolute value of the Pearson correlation coefficient between the observed fMRI time series data). Thus, *M* is simply the unsigned bivariate correlation matrix of the fMRI time series data during a given condition.

Given a brain region *x_i_*, it is natural to ask: which brain region’s activity is *most strongly correlated* with *x_i_*? We say that region *x_j_* is *maximally correlated* to region *x_i_* if |*r* (*x_i_*,*x*_𝑗_) | ≥ |*r* (*x_i_*,*x_j_*) | for all *k* ≠ *i*. That is: *x_j_* is maximally correlated to region *x_i_* if and only if the unsigned correlation coefficient of *x_i_* with *x_j_* is highest among the unsigned correlation coefficients of all regions with *x_i_*, other than *x_i_* itself. We will use the notation *x_i_* → *x_j_* to denote that *x_j_* is maximally correlated to *x_i_*.

There may be “ties” for the region whose correlation coefficient with *x_i_* is highest. For example, we could plausibly have brain regions *x*_1_, *x*_2_, *x*_3_ such that *r* (*x*_1_,*x*_2_) = 0.9 and *r* (*x*_1_,*x*_3_) = 0.9, and such that no other regions’ correlations with *x*_1_ are higher than 0.9. So, several brain regions may be maximally correlated to *x_i_*. Our method will allow such “ties,” although for most sources of real-world data (such as fMRI), “ties” are rare.

As motivated in the Introduction, large-scale networks can be reimagined as terrains, wherein correlation coefficients reflect something like *relative elevation*. The regions in a network can be laid out as points on the topographic map of the terrain in such a way that a marble located at one region *x_i_* rolls toward the region to which *x_i_* is maximally correlated. The path of this marble is presumably curved and winding: if the marble begins its route at region *x*_1_, it may roll first toward a region *x*_2_, then toward a region *x*_3_, then toward a region *x*_4_, rather than *immediately* rolling directly from *x*_1_ toward *x*_4_. This winding behavior arises because *the maximal correlation relation is not transitive*: that is, if *x*_3_ is maximally correlated to *x*_2_, and *x*_2_ is maximally correlated to *x*_1_, it does not follow that *x*_3_ is maximally correlated to *x*_1_.

Consequently, in an idealized situation in which brain regions form a network in which one region *x_i_* plays a central influencing role, but in which there are long “chains” *x_i_* → *x_j_* → *x_k_* → … of maximal correlations starting from *x_i_*, the central role of *x_i_* is not immediately recognizable from the maximal correlation relation, since we do not necessarily have *x_i_* → *x_n_* for all *x_n_* in the network. The problem is the non-transitivity of the relation →. This problem is easily rectified, though. In mathematics, there is a canonical way to construct a transitive relation from any given non-transitive relation. This construction is called the “transitive closure” of the original relation. Given a set *S* and a binary relation < on 𝑆, the transitive closure of < is defined as the binary relation ≺ on *S* given by letting *x* ≺ *y* if and only if there exists a finite sequence 𝑠_1_ < … < 𝑠_𝑛_ in *S* such that 𝑠_1_ = *x* and 𝑠_𝑛_ = 𝑦.

By *transitive maximal correlation*, we mean the transitive closure of the maximal correlation relation. We will write *x_i_* ⇢ *x_j_* to mean that region *x_j_* is transitively maximally correlated to *x_i_*. In concrete terms: *x_i_* ⇢ *x_j_* means that there exists a finite sequence of brain regions *x*_*i*1_,*x*_*i*2_,…,*x*_*i*𝑚_ such that *x_i_* → *x*_*i*1_ → *x*_*i*2_ → … → *x*_*i*𝑚−1_ → *x*_*i*𝑚_ → *x*_𝑗_.

Transitive maximal correlation allows us to define a matrix 𝑇*M*𝐶 (*M*) which is constructed in several steps from the matrix *M*. While the entries in *M* record the bivariate correlations between regions, the entries in 𝑇*M*𝐶 (*M*) are instead determined by which regions are transitively maximally correlated to which other regions. We give the formal definition of *TMC(M)* in Definition 2.1. In section 2.2 we give a simple concrete example of 𝑇*M*𝐶 (*M*).

**Definition 2.1**. Suppose we are given a square matrix *M* of real numbers.

- We put a binary relation on the rows of M as follows. Write *r*_*i*_ for the *i*th row in *M*. Given rows *r*_*i*_,*r*_𝑗_ of *M*, we will write *r*_*i*_ → *r*_𝑗_ if the entry of *M* in row *i* and column 𝑗 is maximal among all entries of *M* in row *i*.
- We write ⇢ for the transitive closure of →. That is, ⇢ is the transitive binary relation on the rows of *M* given by letting *r*_*i*_ ⇢ *r*_𝑗_ if and only if there exists a finite sequence *r*_*i*1_,…,*r*_*i*𝑛_ of rows of *M* such that

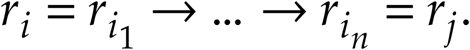

- The *transitive maximal correlation matrix of M*, written 𝑇*M*𝐶 (*M*), is the matrix of real numbers defined as follows: 𝑇*M*𝐶 (*M*) is a square matrix of the same size as *M*, whose entry in row *r* and column 𝑐 is defined to be

- 1 if *x*_*r*_ ⇢ *x*_𝑐_ and furthermore *x_i_* ⇢ *x*_𝑐_ for all regions *x_i_* such that *x*_*r*_ ⇢ *x_i_*,
- 0 otherwise.

To be clear, if we had tried to simplify the definition of 𝑇*M*𝐶 (*M*) by instead writing “The entry of 𝑇*M*𝐶 (*M*) in row *r* and column 𝑐 is defined to be 1 if *x*_*r*_ ⇢ *x*_𝑐_”, then we would have many more 1 entries in the matrix: for example, given a chain *x*_1_ → *x*_2_ →

*x*_3_ → *x*_4_, we would have an entry in row 1 and columns 2 and 3 and 4. Instead, our definition is specifically designed to record the *terminal* entry in any such chain.

Unlike *M*, the matrix 𝑇*M*𝐶 (*M*) is not generally symmetric, because the transitive maximal correlation is not generally symmetric.

In section 2.3 and section 2.5 we give some further explanation of the ideas and intuitions behind the definition of the matrix 𝑇*M*𝐶 (*M*). We conclude this section by pointing out two general properties of the matrix 𝑇*M*𝐶 (*M*):

1. The entry in row *r* and column *r* of 𝑇*M*𝐶 (*M*) is 1 *if and only* if *x*_*r*_ is an attractor region.
2. The trace of 𝑇*M*𝐶 (*M*), i.e., the sum of the diagonal entries in 𝑇*M*𝐶 (*M*), is equal to the number of attractor regions.

**2.2 An example.** Figure 2a depicts a bivariate correlation matrix *M*_1_ derived from hypothetical fMRI time series data.

**Figure 2.**
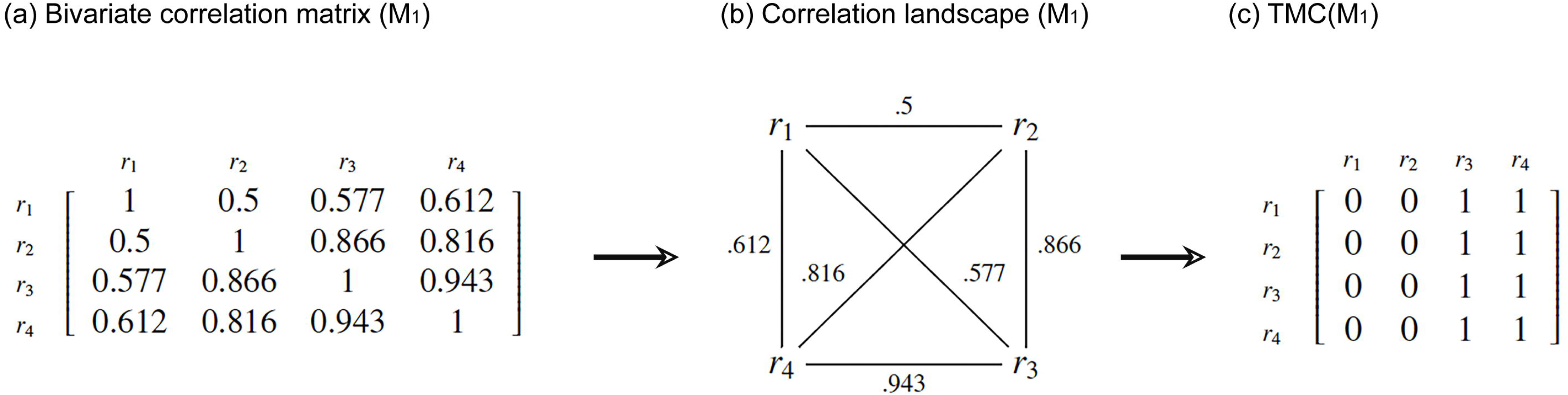
From bivariate correlation matrices to Transitive Maximal Correlations. (a) An example bivariate correlation matrix is depicted for a 4-region space. (b) The data from (a) are represented in an undirected graph or correlation landscape of vertices (regions) and edges (functional distance between regions). For any region *r*_i_, a path can be traversed through the landscape repeating the following example (in this example, i=1): starting at *r*_1_ traverse the edge of highest value to arrive at *r*_4_. From *r*_4_ repeat this algorithm to arrive at *r*_3_. When repeated at *r*_3_ the traversal returns to *r*_4_. Thus, in this example, *r*_3_ and *r*_4_ form an *attractor pair*, and *r*_1_ is *transitively maximally correlated* to each of them. (c) The process from (b) can be represented in a binary Transitive Maximal Correlation matrix. In this matrix, for row *r*_1_ representing that region, we have 1s under columns *r*_3_ and *r*_4_. These values represent the fact that region *r*_1_ is transitively maximally correlated to each of these regions. The remainder of the TMC matrix is filled in by the same method. The resulting TMC matrix in this schematic example is simple, but as can be imagined, in spaces with many regions, the region ◊ attractor pair relations can be more complex.

Figure 2b represents the “correlation landscape” associated to *M*_1_ as a graph, in which each region is a vertex, with edges drawn between all pairs of regions and labelled with the corresponding correlation coefficient. To arrive at the attractor pair for region *r*_1_, we can traverse the landscape beginning at region *r*_1_, and then move along the edge incident to *r*_1_ with the highest label among all edges incident to *r*_1_, i.e., *r*_4_. By repeating this process from *r*_4_, our path eventually loops back and forth between *r*_2_ and *r*_3_, which form an *attractor pair.* Hence the regions *r*_2_ and *r*_3_ are transitively maximally correlated to *r*_1_. This is represented in the corresponding TMC matrix 𝑇*M*𝐶 (*M*_1_) (Figure 2c), where an entry 1 in the row corresponding to *r*_1_ and the columns corresponding to *r*_2_ and *r*_3_ indicates that these regions are transitively maximally correlated to *r*_1_. As 𝑇*M*𝐶 (*M*_1_) shows, *r*_3_ and *r*_4_ are the attractor pair for all the regions in this network.

### 2.3 Directional interpretation of *TMC(M)*

As can be anticipated, transitive maximal correlation, and the transitive maximal correlation matrix 𝑇*M*𝐶 (*M*), are interpretable in terms of *directed* (though *not* causal) relationships between the random variables, an interpretation which can reveal useful structure in the data.

We restate the construction in Figure 2 in graph theoretic terms. Construct a directed graph, with one vertex for each of the brain regions *x*_1_,…,*x_n_*, and in which we draw a directed edge from region *x_i_* to region *x_j_* if and only if *x_i_* → *x*_𝑗_, that is, if and only if *x_j_* is maximally correlated to *x_i_*. Write Γ_*M*_ for this graph. Then *x_j_* is *transitively* maximally correlated to *x_i_*—i.e., *x_i_* ⇢ *x*_𝑗_—if and only if it is possible to start at the vertex *x_i_* and then follow directed edges in Γ_*M*_ and eventually arrive at the vertex *x*_𝑗_.

The entries of “1” in the matrix 𝑇*M*𝐶 (*M*) indicate the *asymptotic behavior* of following directed edges in Γ_*M*_. Generically, there will be no “ties” for maximal correlation. Then, from each vertex *x_i_*, there is *exactly one* directed edge from *x_i_* to some other vertex. In that generic case, if we begin at any vertex *x_i_* in Γ_*M*_ and then follow directed edges, we eventually arrive at a pair of vertices *x*_𝑗_, *x_k_* such that *x_j_* → *x_j_* and *x_j_* → *x*_𝑗_. Continuing to follow edges after that point results in simply moving back and forth between *x_j_* and *x_k_*. As previously noted, such a pair *x*_𝑗_, *x_k_* is an *attractor pair*, and each of *x*_𝑗_, *x_k_* are *attractor vertices*.

𝑇*M*𝐶 (*M*) can be given a graph-theoretic interpretation as follows: we have a 1 in 𝑇*M*𝐶 (*M*) in row *r* and column 𝑐 if and only if *x*_𝑐_ is an attractor vertex such that *x*_*r*_ ⇢ *x*_𝑐_. Otherwise, the entry in row *r* and column 𝑐 in 𝑇*M*𝐶 (*M*) is zero. With such a graph-theoretic interpretation of 𝑇*M*𝐶 (*M*) in place, some matrix-theoretic properties of 𝑇*M*𝐶 (*M*) are clearer. For instance, the diagonal entries are particularly interesting: the entry in row *r* and column *r* of 𝑇*M*𝐶 (*M*) is 1 if and only if *x*_*r*_ is an attractor vertex.

Consequently, the trace of 𝑇*M*𝐶 (*M*), i.e., the sum of the diagonal entries in 𝑇*M*𝐶 (*M*) is equal to the number of attractor vertices.

### 2.4 Distinguishing between 𝐓𝐌𝐂 matrices

Experimentally induced variations in task conditions strongly modulate fMRI signals and network structure [10, 23]. Therefore, TMC matrices are likely to be *condition specific*. In a multi-condition experiment, if *M* is the bivariate correlation matrix of the fMRI data during one experimental condition, and *N* is the bivariate correlation matrix of the fMRI data during another experimental condition, then the differences between the demands evoked by each of the conditions are likely to result in differences between 𝑇*M*𝐶 (*M*) and 𝑇*M*𝐶 (*N*). One of our goals is to demonstrate that in real-world fMRI data these differences are qualitatively meaningful and statistically detectable.

We first consider some idealized “toy” examples in the following section and in Figure 3. Following this, we develop the framework for quantifying condition-driven changes in TMC matrices and in attractor characteristics. Figure 3 depicts network properties that accrue from hypothetical fMRI data where the experiment had three conditions (each row; note that Condition 1 carries over the data from Figure 2). Each row depicts, from left to right, the corresponding bivariate correlation matrix *M*_1_, the transitive maximal correlation matrix 𝑇*M*𝐶 (*M*_1_), and the directed graph Γ_*M*1_ for each condition.

**Figure 3.**
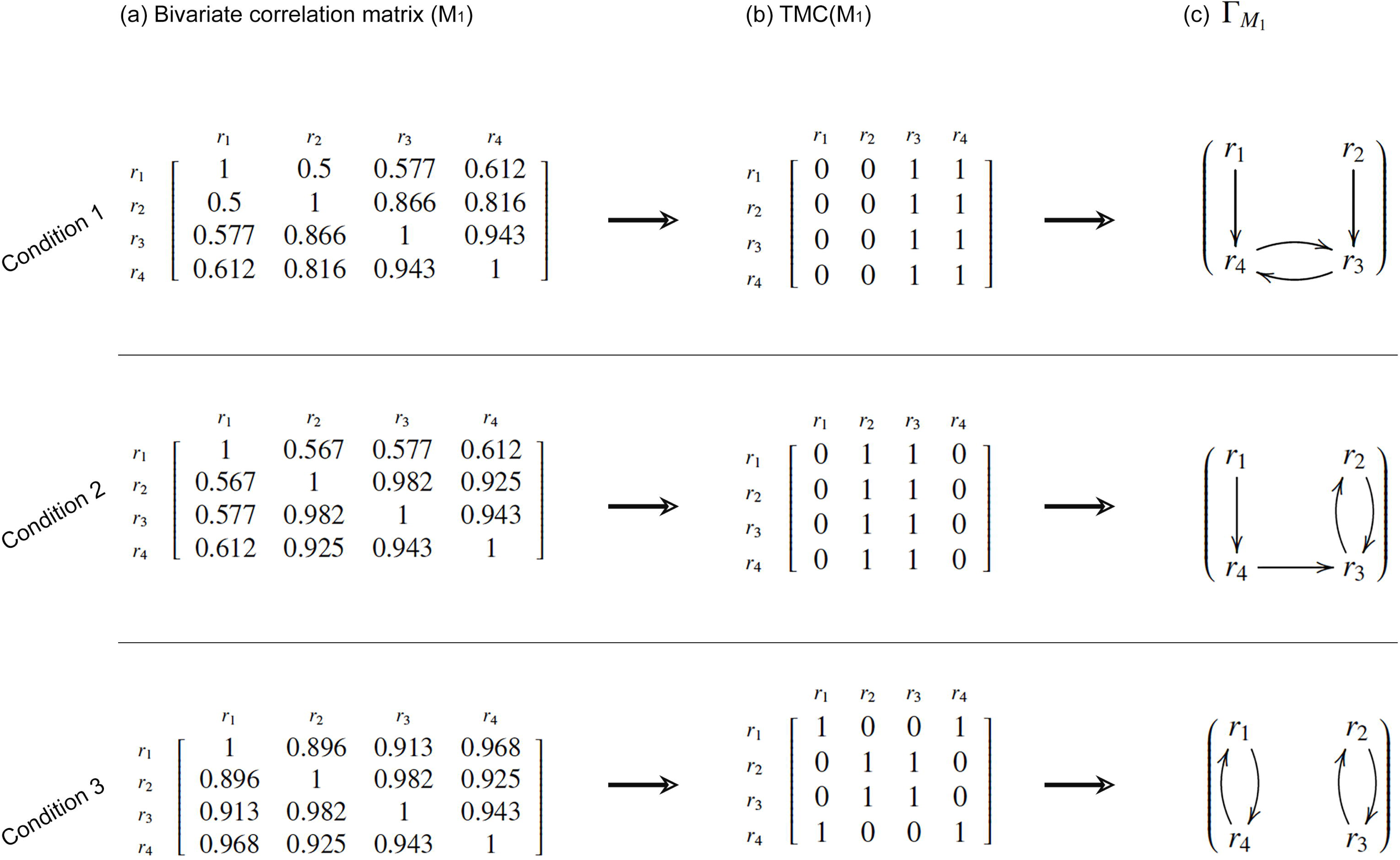
Condition-driven changes in attractor characteristics and representing TMC matrices as “directed” graphs. The schematic example from Figure 2 is carried forward (top row) now representing data from a single condition of any task. The TMC matrix in (b) is effectively a directed graph, depicted in (c). Note that in this condition of the experiment, each of *r*_1_ and *r*_2_ “flow” into attractors r3 and r4. Subsequent rows summarize putative condition-driven perturbations in functional brain interactions. With changes in the bivariate correlation matrices, we also observe substantive changes in the derived TMC matrices, and the corresponding directed graphs. In subsequent sections, we describe how after computing the Frobenius distance between matrices, we can use statistical methods to distinguish between TMC matrices across conditions.

As evident, the second experimental condition evokes a different pattern of network interactions than the first. During the second condition, region *r*_3_ is more strongly correlated to region *r*_2_ than to region *r*_4_. This changes the directed graph Γ_*M*_, compared to Γ_*M*_ : during the first condition, region *r*_3_ “flows into” region *r*_4_, but during the second condition, region *r*_3_ instead “flows into” region *r*_2_. In each of the first and second conditions, precisely two regions are members of attractor pairs. However, a change in task conditions impacts *which* regions are members of attractor pairs. Thus, *r*_4_ is in an attractor pair in the first condition, but not in the second. Conversely, *r*_2_ is in an attractor pair in the second condition, but not in the first.

Condition 3 evokes yet another pattern which now yields two attractor pairs, rather than one, as seen in the transitive maximal correlation matrix 𝑇*M*𝐶 (*M*3) and directed graph *Γ_M3_*. Each task condition evokes qualitative differences in network interactions that are clearly captured across the transitive maximal correlation matrices 𝑇*M*𝐶 (*M*_1_), 𝑇*M*𝐶 (*M*_2_) and 𝑇*M*𝐶 (*M*_3_). *Suitably developed statistical tests applied to the transitive maximal correlation matrices* 𝑇*M*𝐶 (*M*_1_) *and* 𝑇*M*𝐶 (*M*_2_) *can quantify the statistical bases of differences across conditions*.

### 2.5. The dynamics of *TMC* (*M*) across task conditions

As evident from Figure 3, task conditions may induce dynamic changes in 𝑇*M*𝐶 (*M*), where for example, a) as experimental conditions change, some attractor vertices cease to be attractors, while other vertices assume the role of attractors, b) the number of attractor basins might change across conditions, c) regions which previously “flowed” into a given attractor pair instead flow into some other attractor pair. These possibilities are discernable from information contained in the collection of transitive maximal correlation matrices (i.e., 𝑇*M*𝐶 (*M*_1_),𝑇*M*𝐶 (*M*_2_),𝑇*M*𝐶 (*M*_3_),… for the bivariate correlation matrices *M*_1_,*M*_2_,*M*_3_,… of each of the experimental conditions γ_1_,γ_2_,γ_3_ …).

Our next aim is to compare the transitive maximal correlation matrices using a general metric on matrices, which would summarize the differences between the experimental conditions. In mathematics, the *Frobenius distance* is one such metric. It is a standard, simple notion of distance between two matrices of the same size:

**Definition 2.2.** Let *M*,𝑁 be matrices whose entries are real numbers. Suppose that *M* and 𝑁 each have *r* rows and 𝑐 columns. Then the *Frobenius distance from M to* 𝑁, written 𝑑 (*M*,𝑁), is defined as

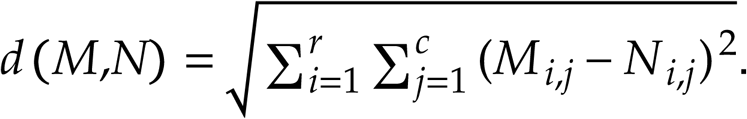

Thus, for any two experimental conditions γ_1_ and γ_2_, we can calculate the Frobenius distance between the two matrices 𝑇*M*𝐶 (*M*_1_) and 𝑇*M*𝐶 (*M*_2_) to arrive at a measure of the aggregate differences in number, size, structure, and composition of attractor basins, evoked by each of γ_1_ and γ_2_.

### 2.6. A statistical test for *TMC* (*M*)

We use the Frobenius distance 𝑑 to define the following test statistic on the two sets of TMC matrices:

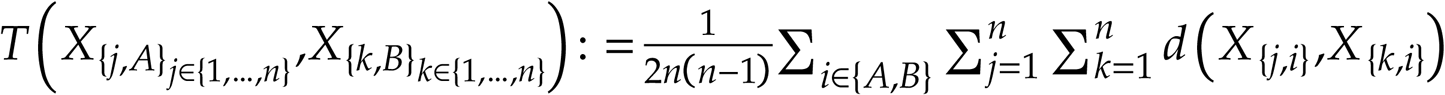

While our interest is to calculate a *p*-value to assess differences in the TMC matrices across conditions, the distribution of our test statistic 𝑇 is not explicitly known and so no distributional assumptions can be made. Therefore, we used a permutation test to carry out this hypothesis testing (the details are given in Appendix 1).

### 2.7. Identifying regions which best distinguish between experimental conditions

A significant *p*-value would indicate that the TMC matrices were significantly different between conditions but would not reveal the regions that were most influential in driving the observed differences between experimental conditions. We addressed this issue as follows: To evaluate the impact a particular region *x* has on the observed test statistic and the corresponding *p*-value, we remove the influence of *x* in the TMC matrix by modifying the matrix where we write zero in all entries in the row *and* column of region *x*. For example, if the region *x* corresponds to the third row in *M*, we replace all entries in row 3 and column 3 of *TMC(M)* with the number zero.

Let *T_x, modified_*be the test statistic, defined in section 2.6, and which is computed from the modified sets of TMC matrices. Let *T_observed_* be the observed test statistic, again as defined in section 2.6, on the *original* (non-modified) TMC matrices. If the difference *T_observed_ -T_x, modified_*is large, then the region *x* plays a pivotal role in the ability of the test statistic *T* to distinguish between TMC matrices. Conversely, if *T_observed_ -T_x, modified_* is near zero, then the region *x* does not greatly contribute to the ability of *T* to distinguish between TMC matrices.

The calculation of the difference *T _observed_ -T_x,modified_*can be carried out for all regions *x*, and the resulting differences can then be ranked by magnitude. This approach allows us to order regions based on how well they distinguished between experimental conditions.

### 2.8 Application to fMRI data and multiple tasks

We applied the TMC framework to fMRI data acquired in two separate domains: a) data collected using an associative learning task known to evoke extensive cross-cerebral and inter-hemispheric interactions [24, 25]. The task has four separate conditions for Encoding, Post-Encoding consolidation, Retrieval and Post-Retrieval consolidation); b) data collected using a basic motor control paradigm with conditions under which participants always responded to the presented visual probe regardless of its color (“sustained response excitation”) or inhibited responses to red probes (33% of trials)(“occasional response inhibition”). MRI parameters, full task descriptions, participant information and details on fMRI processing are provided in Supplementary Materials. Both protocols were approved by Wayne State University’s Institutional Review Board (IRB), and all participants provided consent to participate.

## 3. Results

### 3.1. TMC matrices for the learning and motor tasks

For each of the 39 participants in the learning task, we calculated separate TMC matrices for each of the eight iterations of the four experimental conditions (Encoding, Post-Encoding consolidation, Retrieval, Post-Retrieval consolidation). This yielded 312 TMC (8 × 39) matrices per experimental condition. The averaged matrices in each condition were rendered as heat maps (see Figure 4a for the TMC for Encoding). The 246 cerebral regions in each such heat map are organized by hemisphere (labelled in the figure), and within each hemisphere are listed by their order in the cerebral atlas [26]. Thus, the interhemispheric homologue of the region in the *r*^th^ location lies in the *(246-r)*^th^ location (interhemispheric homologues assume importance in the emerging results).

**Figure 4.**
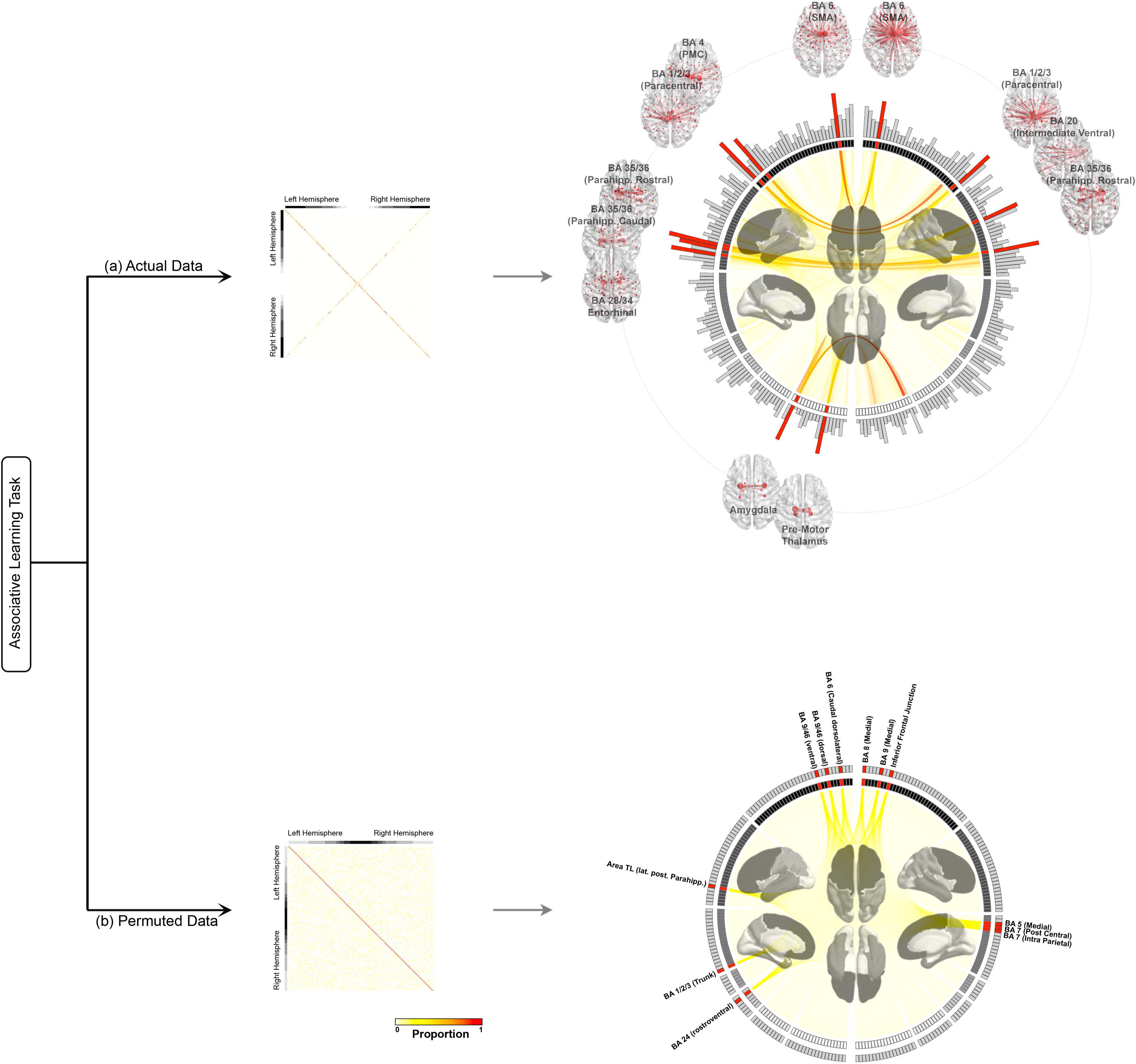
The top sub-figure (a) displays a heat map and chord diagram which are calculated from the actual fMRI data from the Encoding condition of the *Learning task*, while the bottom sub-figure (b) displays a heat map and chord diagram calculated from the fMRI data after its time labels have been randomly permuted (see Appendix B). The heat maps represent the entries in Transitive Maximal Correlation matrices, averaged over iterations of the experiment. The prominence of the main diagonal is an expected phenomenon. This phenomenon occurs because every attractor region is transitively maximally correlated to itself. The prominent skew diagonal, present only in the *actual* data (not the randomly permuted data), is *not* expected. This represents noteworthy structure within the *actual fMRI data* collected in the learning task: *the attractor into which a region flows often includes the interhemispheric homologue of that region*. Thus, in Transitive Maximal Correlation, interhemispheric symmetry can play a more important role than anatomical proximity. This is more lucidly depicted in chord diagrams for the *actual data*. Each chord diagram presents the Transitive Maximal Correlation information from the heat map. The 246 regions (represented by a hash mark in the circle) are organized by hemisphere (right half of the circle is regions from the right hemisphere), and grouped by lobe (see color scheme on the brain template at the center which is carried over to the color scheme on the circle, and in order from top, frontal, temporal, parietal, insular, limbic, visual, and sub-cortical nuclei). The height of the gray bars along the perimeter represents each region’s sensitivity in differentiating between experimental conditions (measured using the method from section 2.7). For both the actual and the permuted fMRI data, each of the twelve maximally distinguishing regions are noted (red hashes). For these, the chord from any region *r*_1_ to each other region *r*_2_ is shaded based on how frequently (i.e., for how many subjects and in how many iterations of the condition) region *r*_1_ is transitively maximally correlated to *r*_2_, i.e., how often *r*1 is in an attractor basin which flows into the attractor *r*2. As noted, it is visually striking that, in the actual fMRI data (top), the maximally distinguishing regions are most often transitively maximally correlated to their interhemispheric homologues. This effect is lost when the data were permuted (bottom). For the actual data, each maximally distinguishing region is also called out in the brain images on the periphery. Here each maximally distinguishing region is connected by a red line to each of the other regions which appears in its attractor basin. The boldness of the red line, and the size of the red dot placed on each of those other regions, varies with the frequency of that other region’s appearance in the attractor basin (larger dots = higher frequency). “SMA” is an abbreviation for supplementary motor area and “PMC” is an abbreviation for primary motor cortex. We provide animated versions of these brain images in the supplementary materials to compare attractor basin composition across conditions.

As previously noted, each cell’s color represents the proportion of the iterations of the condition in which region *c* is an attractor region, and region *r* is transitively maximally correlated to region *c* (i.e., *r* is in the attractor basin which flows into *c*). Therefore, a red cell indicates that, in every iteration of the condition, *c* is an attractor and *r* flows into *c*.

As seen, the main diagonal (from upper-left to lower-right) in Figure 4a is largely red. This indicates that in most iterations of the task condition (in this case Encoding), any region which was an attractor is transitively maximally correlated to itself.

More interestingly, the skew diagonal (from lower-left to upper-right) is also largely red. This effect indicates that on a majority of the task iterations, *each region was also transitively maximally correlated to its interhemispheric homologue*.

Moreover, as is clear from the heat map, most regions are more likely to be transitively maximally correlated to their interhemispheric homologues than with other regions, even anatomically proximate regions. This effect is more clearly elucidated in the adjoining chord diagram which re-represents the Transitive Maximal Correlation information from the heat map. Here, the 246 regions (each represented by a hash mark in the circle) are organized by hemisphere (the right half of the circle shows regions from the right hemisphere) and grouped by lobe (see color scheme on the brain template at the center which is carried over to the color scheme on the circle). Each region’s sensitivity in differentiating between experimental conditions (measured using the method from section 2.7) is indicated in the height of the gray bars along the perimeter (higher bars indicate higher sensitivity). In the figure, a chord from any region *r*_1_ to each of the twelve maximally distinguishing regions (see below) is shaded based on how frequently (i.e., for how many subjects and in how many iterations of the condition) *r*_1_ is transitively maximally correlated to each of those twelve regions (i.e., how often *r*_1_ is in an attractor basin in which each distinguishing region is an attractor). A dark red chord indicates that the regions at the two ends of the chord were members of an attractor pair in many iterations of the task. The twelve maximally distinguishing regions, defined using the methods in section 2.7, are most often transitively maximally correlated to their interhemispheric homologues. These twelve regions are highlighted (red hashes) and depicted in call outs in their approximate location in the brain. In each call out, the red lines connect each region to other regions in its attractor basin.

Further experiments were conducted to investigate whether the inter-hemispheric effect was artifactual or in fact dependent on the canonical ordering of time series in the observed data. First, time indices from the acquired data were randomly permuted in each participant before submitting the permuted data through our TMC pipeline (see Appendix 2 for a full description of these analyses). The resulting heat map in Figure 4b can clearly be differentiated from that in Figure 4a. As seen, the skew-diagonal is no longer present in the randomly permuted data, and the adjoining chord diagram no longer shows a preponderance of chords between inter-hemispheric analogues. The import of these inter-hemispheric effects is discussed in Section 4.4.

Heat maps for data calculated from two conditions of the motor control task are shown in the left-half of Figure 5. Both the main diagonal and skew diagonal are visibly present. However, the inter-hemispheric effect is slightly less prominent than what was observed in the learning task (Figure 4). This effect is further investigated in Section 3.3.

**Figure 5.**
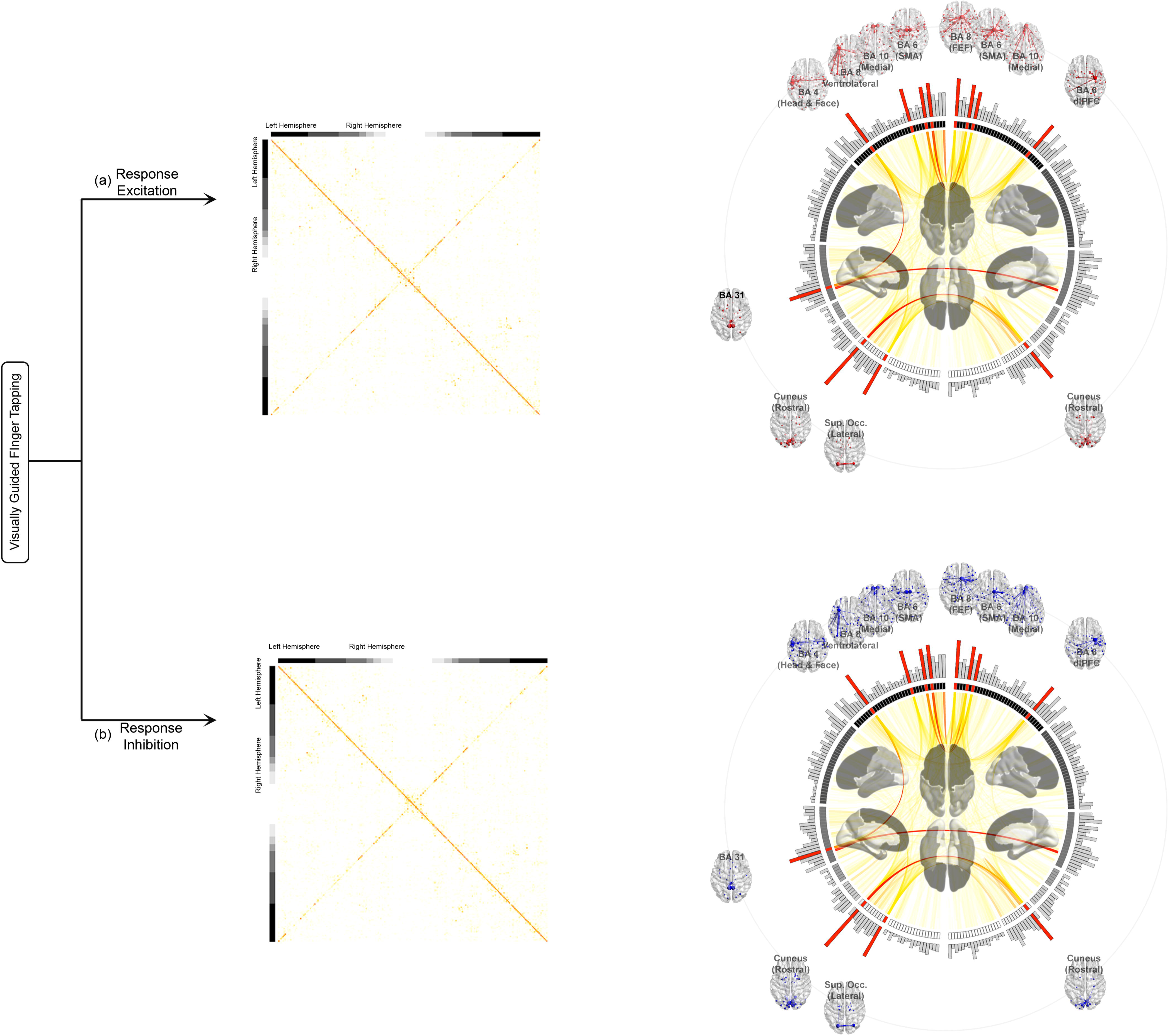
Heat maps, chord diagrams, and anatomical projections were generated from data from the motor (finger-tapping) task in the same way as described for the learning task in Figure 4. The top sub-figure is for the response excitation condition, while the bottom subfigure is for the response inhibition condition. Both instances are derived from the actual fMRI data. As was seen in Figure 4a, the chord diagrams reveal that the strongest connections are predominantly between regions and their interhemispheric homologues. However, as shown in the analysis in Section 3.4, interhemispheric pairing is in fact significantly greater in the learning task. Moreover, for the motor task, the regions which play the most distinguishing roles are different from those observed in the learning task. As in Figure 4, the, maximally distinguishing regions are also called out in the brain images on the periphery (red: response excitation; blue: response inhibition). “SMA” is an abbreviation for supplementary motor area, “FEF” is an abbreviation for frontal eye field, and “dlPFC” is an abbreviation for dorsolateral prefrontal cortex.

### 3.2. Statistical results on condition-specific variation of TMC matrices

We next used the statistical test from section 2.6 to determine whether the TMC matrices differed between experimental conditions. Pairwise comparisons were made between the TMC matrices produced from each of the four conditions in the learning data using the statistical test (with 10k permutations). The *p* values for each comparison are depicted in the heat map in Figure 6(a). They present overwhelming evidence of condition-driven differences in the TMC matrices, indicating that condition-evoked network changes drive changes in the TMC matrices, leading to changes in regional attractors, and the number of attractor basins (see Discussion for further explication). Similar analyses were extended to the data from the more circumscribed motor control task but revealed fewer differences (Figure 6b).

**Figure 6.**
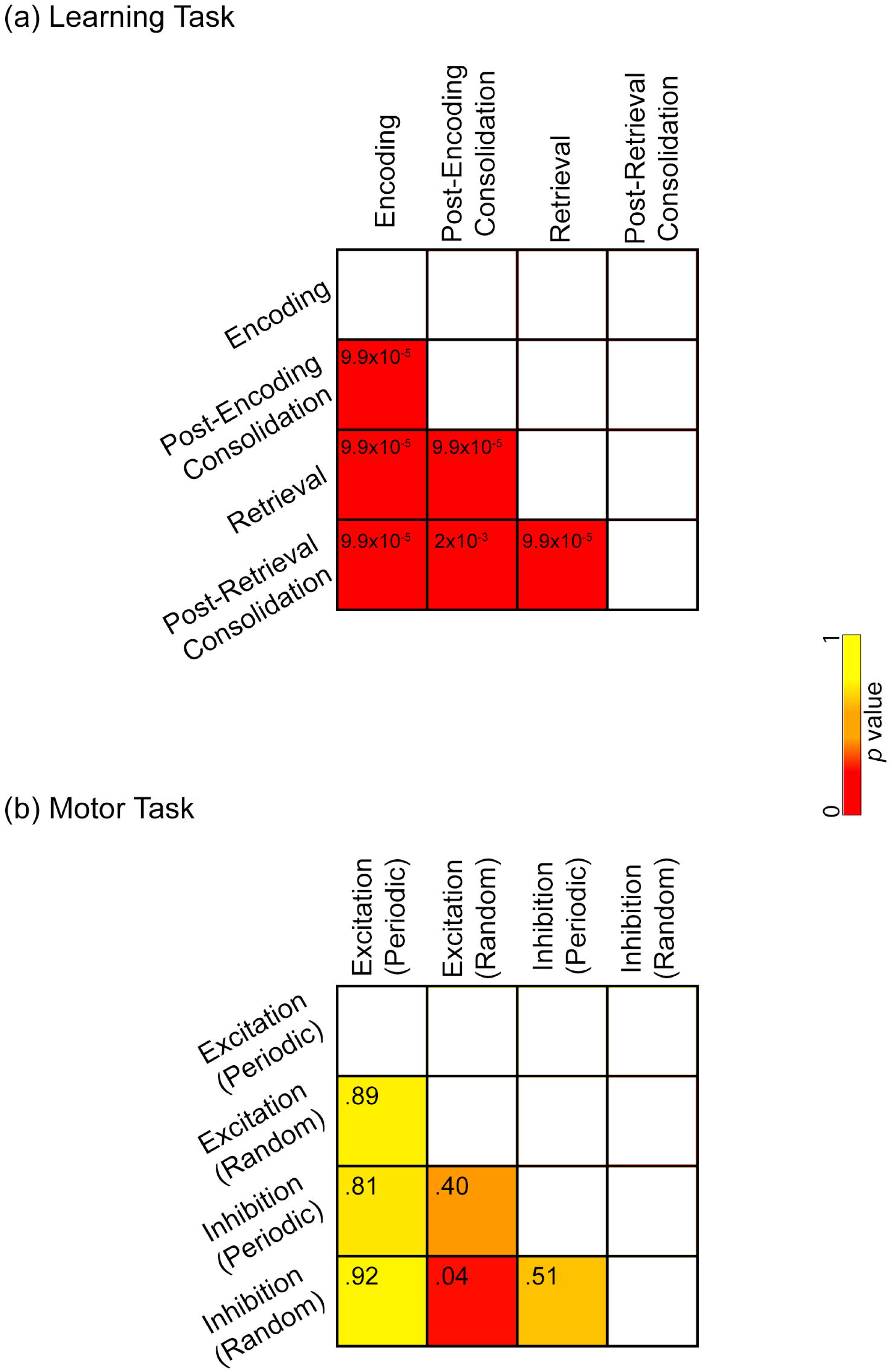
The heat maps provide results from statistical comparisons of TMC matrices from the learning task (a) and the motor task (b). Conditions were compared pairwise using the TMC matrices calculated during each condition and the statistical test from section 2.6 with 10,000 permutations. Each cell in the heat map is shaded from yellow to red based on the p-value for the given comparison (red: lowest p-value; yellow: highest p-value) with observed p-value also reported. For example, the cell in row “Retrieval” and column “Encoding” in (a) corresponds to the statistical test comparing TMC matrices from the Encoding and Retrieval conditions. The composition of attractors and attractor basins for data from the learning task exhibit large condition-specific variations, as is clear from the red cells in (a). However, such condition-specific variation is not present to the same degree for data from the motor task (b). These results indicate that the learning task evokes large-scale and highly contextualized (by condition) network interactions in a way that the motor task does not.

The composition of the average attractor basins for the maximally distinguishing regions is further described by Figures 7 and 8. These heatmaps highlight properties of the attractor basins, including intra-lobular effects, that are not as apparent in the chord diagrams of Figures 4 and 5.

**Figure 7.**
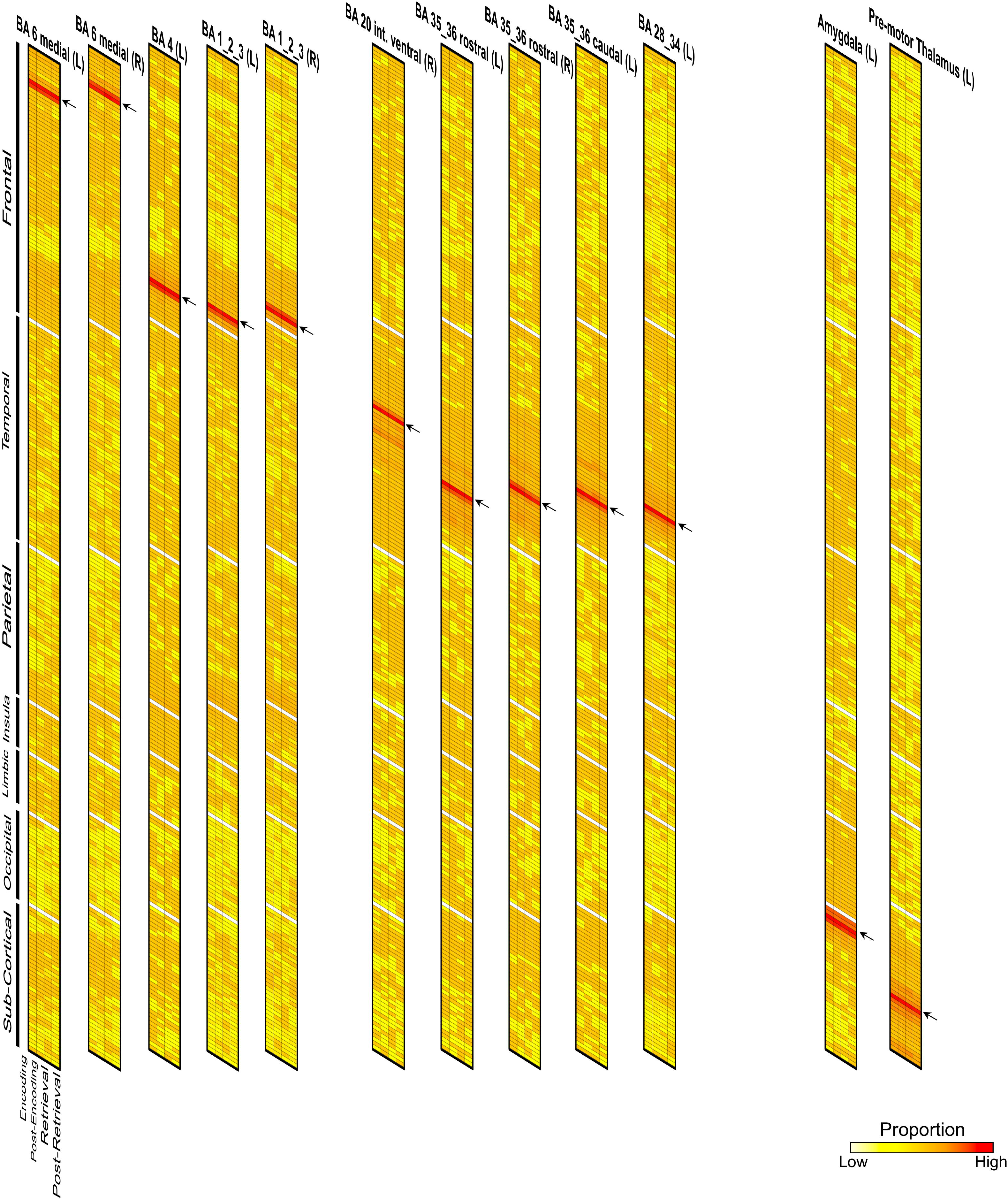
Heat maps are depicted for the twelve maximally distinguishing attractors from the learning task (identified using the method described in Section 2.7). Each column within each heat map represents the average attractor basin composition during the given condition (see condition labels at the bottom). The 246 brain regions are ordered (top to bottom) by lobe. Each arrow points at the row (i.e., region) in the heat map that corresponds to a distinguishing attractor (note that these are the same regions depicted in the periphery of Figure 4a). Each cell’s color is based on the proportion of iterations of the experiment in which the distinguishing region is transitively maximally correlated to the region corresponding to that row (see color bar at the bottom right). Variations in cell colors across columns within a single heat map indicate changes in the average attractor basin composition across conditions. Thus, red cells in successive rows *represent the frequent flow of attractors into their interhemispheric homologues* (and re-represent the interhemispheric chords in Figure 4a). Note, the dense vertical blocks of orange within each attractor’s lobe which indicate that a relatively complex task such as learning reveals strong intra-lobular effects.

**Figure 8.**
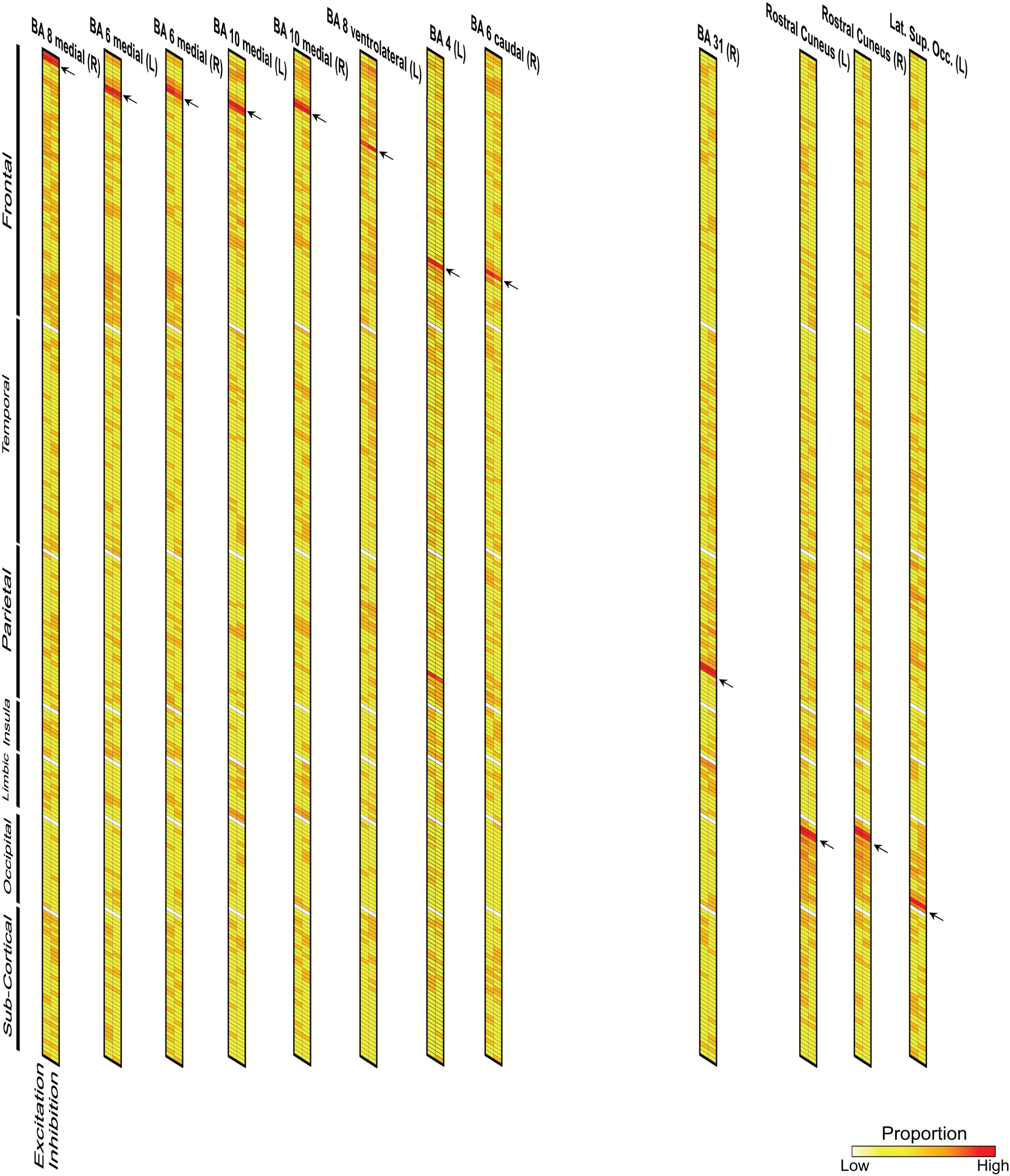
The heat maps for the motor task are depicted in the same format as Figure 7. The differences with data from the learning task are apparent. As seen, yellow cells predominate (compared to what we observed for the learning task).

In the supplementary material, we provide animations that emphasize changes in the anatomical distribution of the attractor basin compositions of each of the maximally distinguishing regions across conditions. The size of the nodes within each animation corresponds to the frequency (across all subjects and epochs), with which each node was present in the attractor basin of the given region (edges are scaled accordingly). Thus, the larger the node, the more often it was present in the given region’s attractor basin. The edges schematically link each node to the attractor. Notably, the thickest edge typically connects interhemispheric homologues. Color encodes experimental condition; red: Encoding, light red: Post-Encoding Consolidation, blue: Retrieval, light blue: Post-Retrieval Consolidation. The largest node in each animation is the given region because it is always present in its own attractor basin. The animation toggles between conditions, making attractor basin changes across conditions easily discernable.

### 3.3. Statistical results on interhemispheric pairing

The interhemispheric phenomenon in transitive maximal correlation is observed in the fMRI data from both the learning and the motor control experiments. However, it appears visually more salient in the data from the learning experiment. To assess whether this observation was quantitatively more pronounced for the learning data, we developed a measure of interhemispheric pairing in the attractor basin for a distinguishing region as follows: For a given condition, let *P_r,D_* be the proportion of iterations of the condition in which a region *r* flows into the attractor basin of a maximally distinguishing region *D*. Let *I* be the set of all regions excluding *D*, and let *I^op^* be the set of regions in the opposite hemisphere of *D*. We define the measure *H(D)* as:

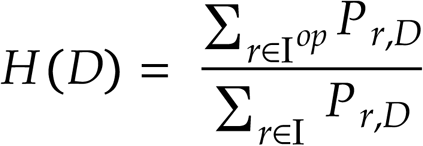

Then *H(D)* directly measures the proportion of interhemispheric pairing in the average attractor basin of *D*. In particular, if *H(D)* is greater than 0.5, then the majority of pairing in the average attractor basin of *D* is interhemispheric.

We computed *H* for the 12 maximally distinguishing regions across the four conditions of interest for the learning experiment, and the two conditions of interest for the motor control experiment (See section 3.2 and Figure 4 for explanation and discussion of “maximally distinguishing regions”). We used the Wilcoxon rank-sum test to test whether there is a significant difference in means between the values of *H* calculated for the two experiments. The result indicated significantly greater interhemispheric pairing in data from the learning compared to the motor control experiment (0.56 vs. 0.40, *p* < 0.0001).

## 4. Discussion

Here, we presented transitive maximal correlation (TMC), a novel method for analyzing fMRI connectivity terrains (Figures 1-2). As we show, TMC is well-suited to identify attractor nodes and basins within any such terrain, and in doing so provides insight on task-induced changes to functional brain network organization. We applied TMC to two independently collected fMRI data sets: a) data co-acquired during a well-established associative learning task known to drive network interactions *on a cross-cerebral basis* [10, 18, 25]; b) data co-acquired during a visually-guided motor control task that drives circumscribed network interactions [21, 27–29]. These were our noteworthy results:

1. In analyzing fMRI data from learning, a) Most attractor pairs were inter-hemispheric homologues (see Figure 4a and Figure 7); b) Different task conditions (Encoding, Post-Encoding Consolidation, Retrieval, Post-Retrieval Consolidation) drove differences in the composition of attractor pairs and basins (see Figure 6a and the animated Supplementary Figures 2-13). Twelve regions significantly distinguished between conditions. These regions were in the paracentral lobule (BA 1/2/3), the motor cortex (BA 4 & 6), the inferior temporal gyrus (BA 20), the entorhinal cortex (BA 28/34), the parahippocampus (BA 35/36), the amygdala, and the thalamus (pre-motor); c) The inter-hemispheric symmetry that we observed for the canonical results (see Figure 4a) was broken when the ordering of time labels was randomly permuted. This indicates that the canonical temporal ordering of data matters (see Figure 4b).
2. We observed a different pattern of results in analyzing fMRI data from the motor control task; a) Here, fewer attractor pairs were inter-hemispheric homologues (see Figure 5 and Figure 8); b) Different conditions during the motor task were less effective at driving differences in attractor node and basin compositions (see Figure 6b); c) regions in the motor cortex were more heavily represented among regions that significantly distinguished between conditions.
3. When comparing results across the two tasks, for the more complex learning task, regions that distinguished between conditions were more likely to be transitively maximally correlated to regions within their lobe (see Figure 7), though this was not the case for the simpler motor control task (see Figure 8).

In the remainder of the paper, we discuss the functional relevance of terms like “attractors” and “attractor basins” and the putative roles that discovered attractors play during tasks such as learning and motor control. We also interpret TMC’s sensitivity to network discovery (e.g., the discovery of inter-hemispheric phenomenon) that may distinguish it from methods like graph theory.

### 4.1. Attractors and attractor basins in TMC

Our definition of attractors is distinct from other definitions. Most notably, in dynamical systems theory, attractors are stable, low-energy states on which the system is likely to converge [30]. Theoretical applications of dynamical systems theory that have modeled fMRI data (notably resting fMRI data) suggest that slow fluctuating (<0.1 Hz) resting state networks emerge as structured noise fluctuations around a stable low firing activity equilibrium state. These arise in the presence of “ghost” multi-stable attractors that together represent the brain’s dynamic repertoire and evidence high signal complexity (i.e., entropy)[31]. These and other investigations [32] have been concerned primarily with the study of non-stationarity and the multi-stability of spontaneously changing brain states. Thus, in a dynamical systems framework, *attractors* are indeed *system states*. By comparison, in the TMC framework, attractors are not system *states* but rather are nodes into which functional paths in a connectivity terrain converge (see Figures 2 and 3, and Methods). As noted, in any bivariate correlation matrix that represents task-evoked system-wide functional connectivity [4, 33], it is possible to derive a functional path that a) begins at any node (i.e., vertex) *n*_i,_ and that after traversing stepwise across each closest connection (edge), b) eventually converges on a pair of nodes, *n*_j_ and *n*_k_ from which the formed path cannot escape. Here, *n*_j_ and *n*_k_ form the attractor pair into which *n*_i_ converges, and the set of all such *n*_i_ is the attractor basin for that pair. In effect, TMC *partitions any large network into multiple loci (of attractor basins)* where each such locus represents the unique smallest sub-network that contains each of its members (assuming Axiom 1 and Axiom 2 in the Introduction). These partitions or sub-network configurations are derived from purely local operations and are distinguishable from other established techniques like graph theory.

### 4.2. Comparison to graph theory

How is TMC is related to, or unique from graph theoretic measures including path-based measures such as characteristic path length and node-based measures such as betweenness centrality [10, 34]? The characteristic path length of a network is the average shortest path between all pairs of nodes and is a single quantity that measures a global property of the network [35, 36]. By comparison, while betweenness centrality is sensitive to global network properties [34], it specifically quantifies how often a node lies on the shortest path between all pairs of nodes in a network [9, 14]. Both measures respectively provide information about the integrative capacity of the network or the integrative role of individual nodes in the network.

However, they are not suited for partitioning networks based on *local functional* properties (though betweenness centrality can be used to identify hub-like structures in brain networks) [37, 38].

It is straightforward to demonstrate that TMC differs from betweenness centrality.

Consider the symmetric correlation matrix in Figure 9.

**Figure 9.**
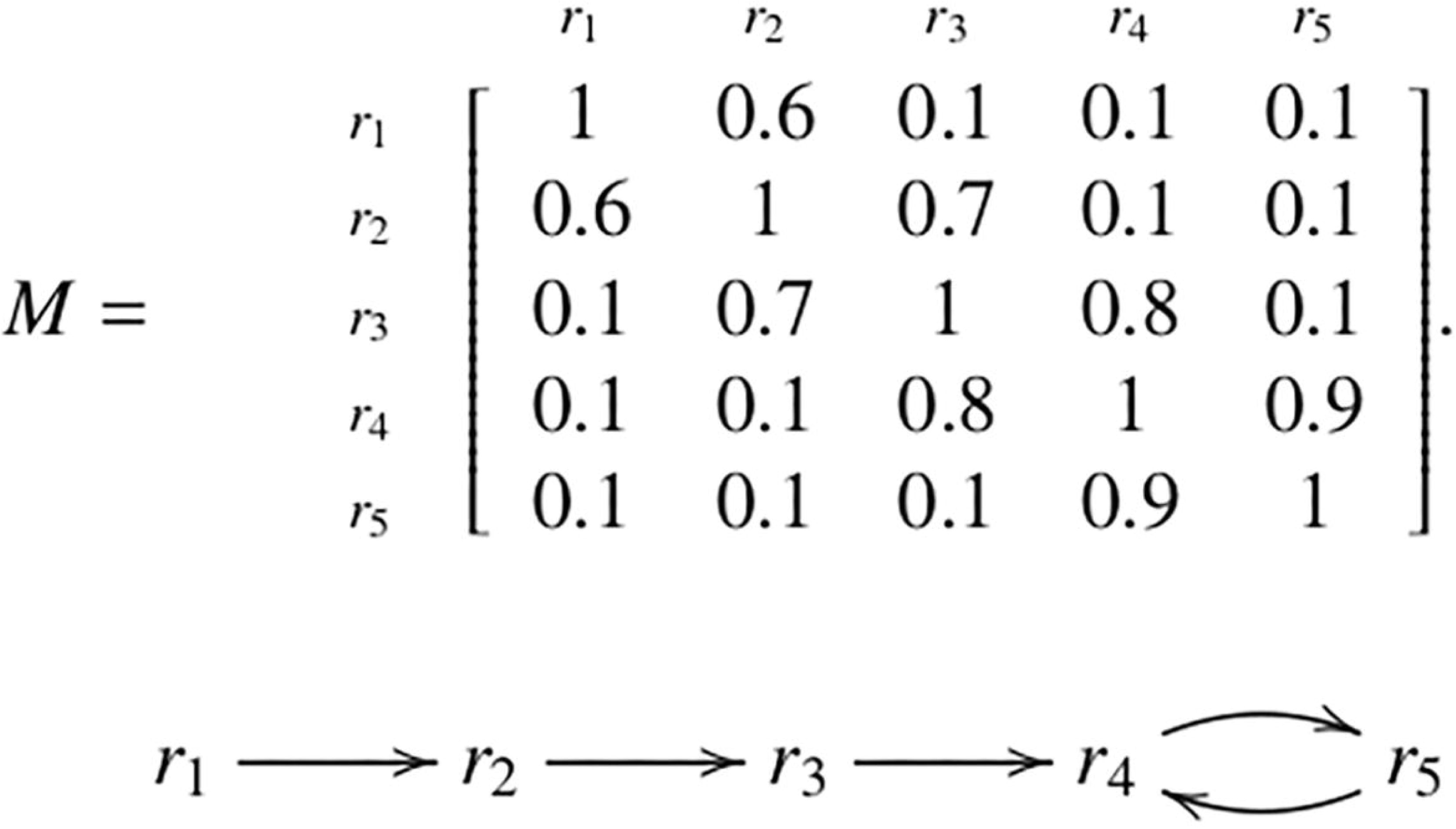
In this toy example, the given bivariate symmetric correlation matrix M yields the maximal correlation graph pictured below **M**. The regions *r*4 and *r*5 are the attractor regions. Nevertheless, the betweenness centrality is highest at region *r*_3_. This simple example demonstrates that attractor regions are not, in general, the same as regions of highest betweenness centrality. Consequently, the techniques of Transitive Maximal Correlation do not detect the same features as the methods of betweenness centrality.

Here, *r_3_* has the highest betweenness centrality of 8, as the shortest weighted path from *r_i_* to *r_j_* passes through *r_3_* if *i*=1, 2 and j= 4, 5, or if i=4, 5 and j=1, 2. A similar calculation shows that the betweenness centrality of *r_2_* and of *r_4_* is equal to 6, and the betweenness centrality of *r_1_*and of *r_5_* is equal to zero. However, this does not constitute a partitioning of the network but rather summarizes aspects of global network organization at the level of each individual node.

By comparison, the maximal correlation graph Γ*_M_* identifies *r_4_* and *r_5_* as the attractor regions. This example shows that a region can have very high integrative value based on global metrics like betweenness centrality, but local measures like TMC show that the same region is not part of an attractor pair. Conversely, a region (such as *r_5_*) might have low betweenness centrality (which for *r_5_* is zero) but be part of an attractor pair.

We further advanced this example by comparing betweenness centrality and TMC in our own fMRI data. These analyses showed that there is no principled relationship between attractor status and betweenness centrality. Figure 10 shows the density plots for attractors and non-attractors for each of the a) learning and b) motor control data (Appendix 3 provides details on statistical tests used to assess differences between the mean betweenness centrality ranks of attractors against ranks of non-attractors). For the learning data, attractor regions had significantly higher betweenness centrality than non-attractors (*p*<0.0001). However, this effect was not observed for the motor control data (*p*>.25). Investigations on the permuted data (Appendix 2) further reinforced a lack of any principled relationship. More in-depth investigations of this issue exceed our paper’s scope, but we suspect that any such investigations will confirm the complementary relationship between TMC and graph theoretic measures like betweenness centrality.

**Figure 10.**
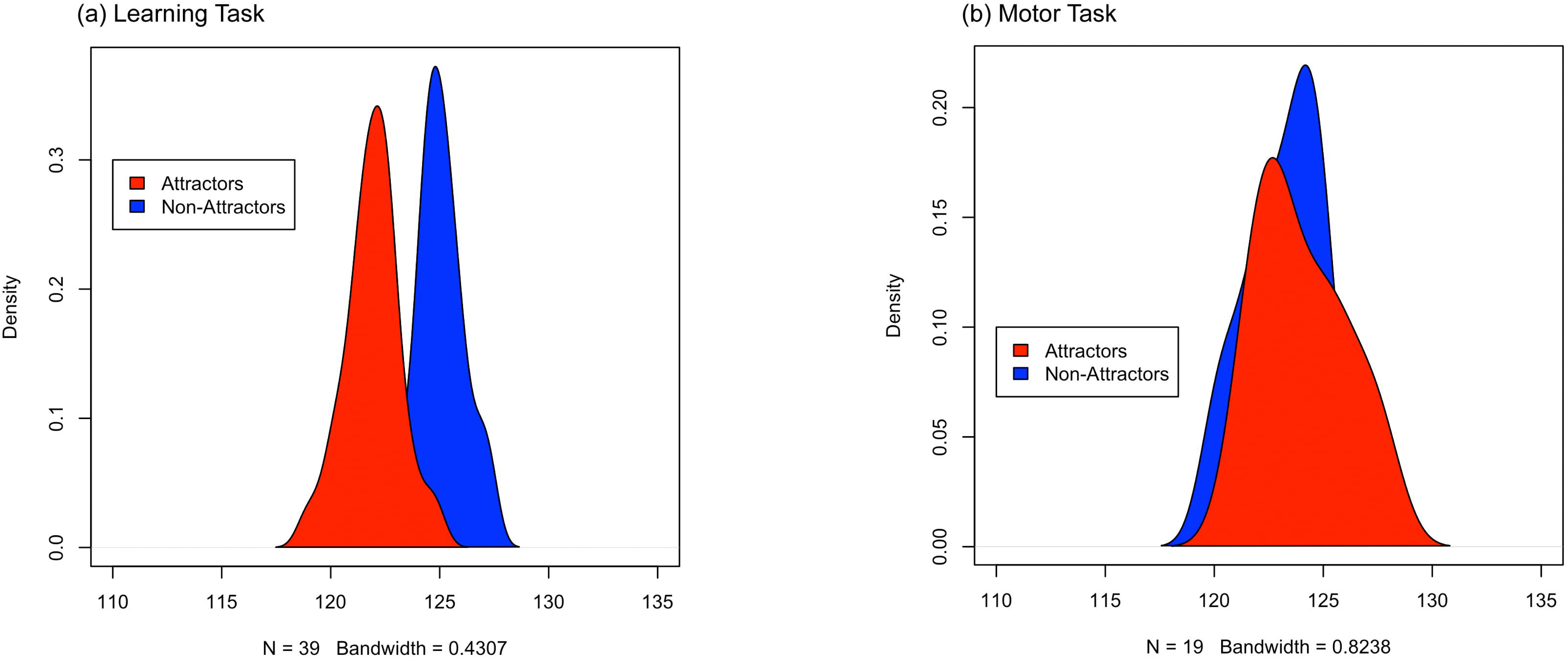
Density plots derived from histograms of subject means for the betweenness centrality (BC) rank of attractors and non-attractors in the learning task (a) and motor task (b) as calculated in Appendix 3. The x-axis represents the BC rank, while the y-axis represents the probability density. The distributions for attractors are colored in red and the distributions for non-attractors are colored in blue. The clear visual difference in mean BC ranks between attractors and non-attractors in (a) is supported by a significant Wilcoxon rank-sum test result. In contrast, (b) shows substantial overlap between the distributions, consistent with an insignificant result from the Wilcoxon rank-sum test. These results demonstrate that attractors, on average, have higher integrative value than non-attractors in the learning task, while no such phenomenon is present in the motor task. The relationship between Transitive Maximal Correlation and betweenness centrality is task-dependent, suggesting that they capture different (perhaps complementary) properties in brain networks.

### 4.3. Task-relevant identities of the Maximally Distinguishing Regions

The procedure from section 2.7 does not reveal whether a region’s status as one that maximally distinguishes between conditions is driven by a) changes in its attractor status or b) changes in its attractor basin composition. We addressed this question by applying a modified version of the test from section 2.6.

Beginning with the full set of TMC matrices for the learning data, we restrict the test from section 2.6 to the 1x1 submatrices given by the row and column of a given maximally distinguishing region. Recall that this cell of the TMC matrix contains a one if a region is an attractor and a zero otherwise. This modified test is therefore only sensitive to a region’s attractor status. Six tests with 10k permutations were performed for each of the 12 maximally distinguishing regions, one for each pairwise comparison. The FDR-corrected *p*-values (Supplementary Figure 1) showed that none of the tests were significant (α < 0.01). These findings suggest that changes in attractor basin composition, rather than changes in attractor status drive the maximally distinguishing role of regions (the changes in attractor basins are observable in the animations that form Supplementary Figures 2-13). Thus, the attractors appear to be regions that play stable and well-identifiable functional roles throughout the entire task, but whose attractor basin (i.e., sub-network) compositions change. This appeared to be generally true for both the learning (Figures 4,7) and the motor control data (Figures 5,8).

For example, during learning, these regions were in the rostral and caudal parahippocampus (BA 35/36), the entorhinal cortex (BA 28/34) and the ventral temporal lobe (BA 20). The entorhinal cortex and the parahippocampus are key nodes in a cortical memory network hub the dynamics of which underpin associative learning [39–41]. The basolateral amygdala modulates hippocampus-dependent memory consolidation by impacting synaptic strengths in hippocampal structures [42]. The ventral temporal cortex processes visual shape information and is a central element of the ventral visual pathway [43–46], supplementing the functions of the medial temporal lobe in learning and memory [46]. Areas of the somatosensory cortex (BA 1/2/3) are also known to supplement long term memory formation through involvement in visual short-term memory maintenance [47]. Finally, as an active gateway to the cortex, the thalamus is involved in retaining short-term information related to spatial cognition and spatial learning [48, 49]. By comparison, the distinguishing regions from the motor control data showed a) less diversity of function (Figure 5) and b) were more closely associated with functions related to finger movements, movement initiation and response inhibition [21, 50, 51].

### 4.4. Interhemispheric phenomenon

An unexpected finding was that many of the distinguishing regions were transitively maximally correlated to their interhemispheric homologues (notably for the learning task, see the heatmap and chord diagram in Figure 4a). This indicates that many attractor basins were formed from inter-hemispheric homologues. This finding obviously indicates that the anatomical proximity between pairs of nodes was not a predictor of the likelihood that they would form attractor pairs. In TMC, attractor basins are derived entirely from the iterative application of *local operations on each node* (see Methods) and therefore might *a priori* be expected to be *anatomically* proximate. Therefore, the inter-hemispheric effects are somewhat unexpected. However, this evidence may reveal TMC’s ability to capture global principles of sub-network organization. While anatomical proximity is *one* organizing principle for how the brain’s functional sub-networks are formed, this principle is more evident during tasks that rely on greater regional cooperation over global synchrony [52, 53]. In general, however, dependence on anatomical proximity places an arbitrary constraint on the brain’s ability to implement functional diversity. Indeed, the transition from local to distributed function is a feature of brain development [54] and the brain’s adaptiveness stems from its ability to dynamically form (and dissipate) local *and* global brain networks [55]. This may also be explained by the fact that exuberant inter-hemispheric connectivity is a general feature of mammalian brains [56], and at least in humans, long distance inter-hemispheric synchronization may be central to the maintenance of conscious awareness [57, 58]. Thus, for tasks like learning that have a global footprint, TMC may be sensitive in identifying sub-networks that are formed around inter-hemispheric attractor basins. This promises to be one more unique role for the method in network neuroscience.

### 4.5. Conclusions and Limitations

Network neuroscience offers analytical approaches that might breach the barriers to our understanding of brain function [59, 60]. Amongst the methods for network neuroscience that take connectivity terrains as their starting point, TMC offers a novel approach to revealing organizational features in brain networks. The outputs of TMC, in the form of attractors and attractor basins, identify a) influential brain regions within the network (attractor pairs) and b) partitions brain networks into functionally proximate sub-networks (attractor basins). Our investigation provided some of the mathematical foundations for TMC, statistical approaches that can be used to distinguish between TMC matrices formed from different task conditions, and applications of this framework to two unique fMRI datasets. Our results are compelling enough to suggest that the TMC construction is broadly sensitive to the demands of the task (whether more global as in the learning task, or more circumscribed as in the motor control task). TMC inherits many of the limitations that are inherent in working with correlational data, including hidden drivers of signals, noise and other fMRI signal artefacts [4, 61, 62]. However, we suggest that our theoretical and empirical demonstrations establish TMC as a unique and high impact method for functional network discovery. The method can readily be applied for multifarious applications, including single-subject analyses, the characterization of resting state networks, and the identification of aberrant functional organization in neuropsychiatric conditions. We hope that the tool is seen as an important method in the arsenal used by network neuroscientists.

## 5. Appendix 1: the permutation test for hypothesis testing using TMC

We defined a permutation test for hypothesis testing based on the test statistic *T* defined in section 2.6. Broadly, a permutation test uses a null distribution that is empirically generated by permuting labels and recomputing the test statistic many times over. Let *T(L_observed_)* be the value of the test statistic on the observed labels *L_observed_*. We obtained the probability of obtaining the observed test statistic under the null hypothesis, i.e. a p-value, by taking the proportion α of labelings *L* such that *T(L)* ≤*T(L_observed_)*. In our application of interest, care must be taken when generating the empirical null distribution. For instance, if the labels of experimental condition *A* and experimental condition *B* are permuted without restriction, one can imagine that the resulting test statistic may be extreme as a result of confounding factors.

For instance, consider the permutation of labels where all TMC matrices originating from the first two runs of the experiment are labelled *A* and all the TMC matrices originating from the last two iterations of the experiment are labelled *B*. The test statistic associated with such a permutation could potentially detect a difference between the two groups not arising from a difference in the experimental conditions. To guard against the unpredictable effect of intersubject variability, only permutations that result in an equal representation of subjects and runs of the experiment in each group are permissible. Details about permuting labels under such restrictions can be found in [63].

Once a *p*-value is obtained, it is compared to an *a priori* chosen threshold to determine whether or not to reject the null hypothesis. Rejecting the null hypothesis would indicate evidence of a significant difference between the two sets of TMC matrices, and therefore a meaningful difference of the system properties of transitive maximal correlation of the brain during experimental condition *A* and experimental condition *B*.

## 6. Appendix 2: Randomization experiments

This section aims to determine which properties of the TMC matrices in our results depend on the temporal structure of the experiment (i.e. the actual time and condition labels attached to each time point). We accomplished this by disassembling the temporal structure of the data via randomization before computing the corresponding TMC matrices. The fMRI data in the study described in section 3.1 was collected at 288 time indices for each subject, so we carried out the following temporal randomization experiment:

- For each of the 246 regions, we freely permuted the vector (1,2,…,288) and reordered each region’s fMRI time series based on the resulting permuted vector. Using the actual condition and iteration labels for each time index, we then computed the TMC matrix corresponding to each condition, iteration, and subject.
- We then applied the statistical test of section 2.6 to compare sets of TMC matrices based on the actual condition labels.
- Steps 1 through 3 were repeated 1000 times.
- We then averaged the TMC matrices generated for the actual condition labels across all subjects.

The null hypothesis was rejected in 1 out of the 6000 tests from Step 4 after applying a multiple testing correction using the False Discovery Rate. The average TMC matrices generated for each condition (see the bottom subfigure of Figure 4) display a prominent main diagonal, for the reasons described in Section 3.3, but without a prominent skew-diagonal. The non-prominence of the skew-diagonal indicates that the interhemispheric pairing phenomenon, which is clearly present in the non-permuted fMRI data, is lost after the random permutation.

## 7. Appendix 3: Comparison to betweenness centrality

Betweenness centrality is a graph-theoretic measure commonly used in the analysis of functional connectivity data. It is a measure of the number of shortest paths between pairs of nodes that a given node lies on. This section assesses the relationship between our attractors and betweenness centrality (BC). Specifically, we aimed to determine if the mean BC rank for attractors differs from that of non-attractors.

We calculated the attractor status and BC rank of regions using the bivariate correlation matrices (246 x 246) produced from the associative learning experiment (39 subjects x 4 conditions x 8 iterations = 1248 matrices) and finger tapping experiment (19 subjects x 4 conditions x 2 iterations = 152 matrices). BC scores were calculated considering *stronger* functional connectivity as *shorter* edge length. Figure 10 displays the distribution of subject means for the BC rank of attractors and non-attractors for the learning data and finger tapping data. We compared the sets of subject means using the Wilcoxon rank-sum test. While the test did not detect a significant difference in mean BC ranks between non-attractors and attractors for data from the motor control task (124.0 vs 123.1 *p*=0.275), a significant difference was detected for data from the learning task (121.7 vs 125.3, *p*<0.0001). These results indicate that, in the learning data, attractor regions tend to have a higher integrative value as measured by betweenness centrality than non-attractor regions.

We repeated the above analysis with randomized data to further probe the relationship between BC and our attractors. Specifically, we randomly permuted off-diagonal elements of the bivariate correlation matrices from the learning experiment before calculating BC ranks, identifying attractors and non-attractors, and comparing subject mean BC ranks for attractors and non-attractors using the Wilcoxon Rank-Sum test. We carried out 100 permutations of this procedure and found a significant difference in subject mean BC rank between attractors and non-attractors in only 4 percent of permutations, revealing that attractor status and BC rank are generally not related in the randomized data. The resulting density plots are given in Figure 10.

## 8. Appendix 4: Proof of the theorem on minimal subnetworks

In this appendix, we present the proof of the theorem stated in the introduction:

**Theorem**. Given a member r of S, there is a unique smallest sub-network in S containing r. This smallest sub-network is precisely the attractor basin containing r.

The proof is not difficult, and could be worked out by an interested reader, but for the sake of completeness, we present it here.

**Proof of theorem.** Given a collection of subsets of *S* to be regarded as sub-networks, we will write *x ∼ y* if *x,y* are members of *S* and some sub-network in *S* contains both *x* and *y*. For a given such collection of subsets of *S*, it is easy to check that the relation *∼* on *S* is symmetric, transitive, and reflexive, i.e., it is an equivalence relation. The second axiom, above, ensures that if *x* is maximally correlated to *y*, then *x ∼ y*. From our construction of transitive maximal correlation as the transitive closure of maximal correlation, and from the observation that *∼* is transitive, it follows that whenever *x* is transitively maximally correlated to *y*, we have *x ∼ y*.

Now suppose that *x* and *y* are in the same attractor basin. Then there exists an attractor *z* such that *x* and y are each is transitively maximally correlated to *z*. We have *x ∼ z* and *y ∼ z.* Since the relation ∼ is symmetric, we have *z ∼ y,* hence *x ∼ y* by transitivity.

Hence, if *x* and *y* are in the same attractor basin, then every sub-network that contains either of *x* or *y* contains both of them. Hence the smallest sub-network containing a single member of S also contains that member’s entire attractor basin. Q.E.D.

## 9. Supplementary Materials

### Participants, Tasks and MRI acquisition

All protocols were approved by Wayne State University’s Institutional Review Board (IRB). All participants provided written consent to participate in the protocols were compensated for their participation. They were free of past or present Axis-I psychopathology (assessed by a clinical psychologist using the DSM-5)[64] and were screened to exclude any significant past/current medical and/or neurological illness (e.g., hypertension, thyroid disease, diabetes, asthma requiring prophylaxis, seizures, or significant head injury with loss of consciousness). fMRI data were collected from these 58 healthy participants in two separate cohorts for learning (n=39, 10 females, mean age=28.2 yrs.) and for motor control (n=19, 5 females, mean age=18.36 yrs.). <u>Tasks</u>. *Learning*: Learning was evoked using an established paired-associative learning paradigm that has been previously described [10, 17, 34, 36, 65, 66]. The task required participants to learn associations between nine equi-familiar objects [67] and their assigned grid locations. The eight task iterations cycled between epochs (27 s) for Encoding, Post-Encoding Consolidation, Retrieval, and Post-Retrieval Consolidation. During Encoding, objects were presented in their assigned grid location for naming (3 s/associated pair). Following a 27 s passive Post-Encoding Consolidation period, learning proficiency was tested in a Retrieval epoch. Here, each of the nine grid locations were highlighted (in random order) and participants were required to recall and name the associated object (or indicate “no”).

The entire paradigm lasted 864 s. Though hippocampal centric, associative learning has a cross-cerebral signature and is known to drive connectivity between diverse regions across the cerebral and sub-cortical nuclei [34, 36, 68, 69]. *Motor Control*: The paradigm is a variant of previously deployed tasks for visually-guide motor control [21, 22, 27, 29] where such control is evoked by manipulating the relationship between the presented stimulus (Red or Green probes) and the expected response. During epochs (32 s) for sustained response excitation, participants were instructed to respond to the presented probe regardless of its color. During epochs (32 s) for occasional response inhibition, participants were instructed to respond only to green probes (67%). Rest epochs (16 s) were interspersed throughout the task. The entire paradigm lasted 384 s. Motor control has a comparatively circumscribed cross-cerebral signature, and is known to primarily drive interactions between the pre- and supplementary motor areas, the primary motor cortex, the dorsal anterior cingulate and the basal ganglia [21, 50, 70].

#### MRI acquisition

All MRI data were acquired at the WSU MR Center using a 3T Siemens Verio scanner using a 32-channel volume head coil. *Associative Learning*: fMRI data were acquired using a multiband gradient EPI sequence with the following parameters: TR = 3 s, TE = 24.6 s, multiband factor = 3, FOV = 192 × 192 mm^2^, matrix = 96 × 96, 64 axial slices, resolution = 2 mm^3^). In addition, we also collected T1-weighted MRI images were collected for normalization and co-registration with the EPI scan (3D magnetization-prepared rapid gradient-echo sequence, TR = 2,150 ms, TE = 3.5 ms, TI = 1,100 ms, flip angle = 8°, FOV = 256 × 256 × 160 mm^3^, 160 axial slices, resolution = 1 mm3). *Motor Control*: fMRI data were acquired using the following parameters: TR: 2600 ms, echo time; TE: 29 ms; matrix dimensions: 128 × 128; voxel dimensions: 2 mm^3^; FOV = 256 × 256 mm^2^; 36 axial slices; pixel resolution: 3.75×3.75×4.0 mm^3^. A T_1_-weighted structural image was acquired using a 3D Magnetization Prepared Rapid Gradient Echo (MPRAGE) sequence with the following parameters: TR: 2,200 ms; TI: 778 ms; TE: 3 ms; FOV: 256 mm^3^; matrix dimensions 256 x 256; flip angle: 13°; 256 axial slices of 1.0mm thickness; pixel resolution: 1 mm^3^. In both acquisitions, axial slices were positioned parallel to the anterior commissure/posterior commissure (AC-PC) line and provided comprehensive cerebral (but not cerebellar) coverage.

### MRI Processing

MRI data were processed using standard temporal (slice-time correction) and spatial preprocessing methods in SPM 12. The EPI images were aligned to the AC-PC line, realigned to a reference image in the sequence to correct for head movement, and co-registered to the high-resolution T1 anatomical image. The EPI images were normalized to stereotactic space by applying the deformations from normalizing the high-resolution T1 image. A low-pass filter (128 s) was used to remove low frequency components associated with respiratory and cardiac rhythms. At the first level, boxcar stimulus functions were convolved with a canonical hemodynamic response function modeled epochs (in each of the tasks) as regressors of interest with the six motion parameters (3 for translation and 3 for rotation) from the co-registration modeled as covariates of no interest. The images were resliced (2 mm^3^), and a spatial filter was applied (4 mm FWHM). Images with more than 4 mm of movement (<1% of all images) were excluded from analyses. This processing pipeline emulates previously published work in clinical and non-clinical populations [18, 25, 34, 36, 66, 71, 72]. From the pre-processed fMRI data, time series were extracted from each of 246 regions in the Brainnetome cerebral atlas [26, 34, 36] (chosen because the parcellation scheme is based on multi-modal imaging data acquired in the same group of participants). Next, bivariate zero-lag correlations (undirected functional connectivity) [4] were estimated between all 30,135 pairs of regions (^246^C_2_). For the associative learning paradigm, these were estimated for each participant in each iteration (eight) of each of the conditions (four in total). For the motor control task, these were estimated in each of the two task conditions.

## Supplementary Figure Captions

Supplementary Figure 1. We provide results from testing whether a region’s status as a discriminator between conditions was driven by changes in attractor status *or* changes in basin composition. Beginning with the full set of TMC matrices (for the learning data), we restricted the test (from section 2.6) to the 1x1 submatrices given by the row and column of a given maximally distinguishing region. This cell of the TMC matrix contains a one if a region is an attractor and a zero otherwise. This modified test is therefore sensitive to a region’s attractor status. Six tests with 10k permutations were performed for each of the 12 maximally distinguishing regions, one for each pairwise comparison. The FDR-corrected *p*-values (Supplementary Figure 1) showed that none of the tests were significant (α < 0.01). Each bar height represents the p-value for the pairwise test for that region. As seen, no bar passes the significance threshold (red line). These findings suggest that changes in attractor basin composition, rather than changes in attractor status drive the maximally distinguishing role of regions. These changes in attractor basins are observable in the animations that form Supplementary Figures 2-13.

Supplementary Figure 2. The animation represents the average attractor basin composition for (BA 1/2/3 L). The size of the nodes corresponds to the frequency (across all subjects and epochs), with which each node was present in the attractor basin of BA 1/2/3 L (edges are scaled accordingly). Thus, the larger the node, the more often it was present in BA 1/2/3’s attractor basin. The edges simply schematically link each node to the attractor (i.e., they do not represent the path from the attractor to each node). The edge color encodes experimental condition; red: Encoding, light red: Post-Encoding Consolidation, blue: Retrieval, light blue: Post-Retrieval Consolidation. BA 1/2/3 is the largest because it is always present in its own attractor basin. This scheme is carried forward for all subsequent animations (Supplementary Figures 3-13).

