## Supplementary figures and images for "A novel mathematical construction for identifying attractors from task-driven fMRI data"

### Supplementary Figure 1

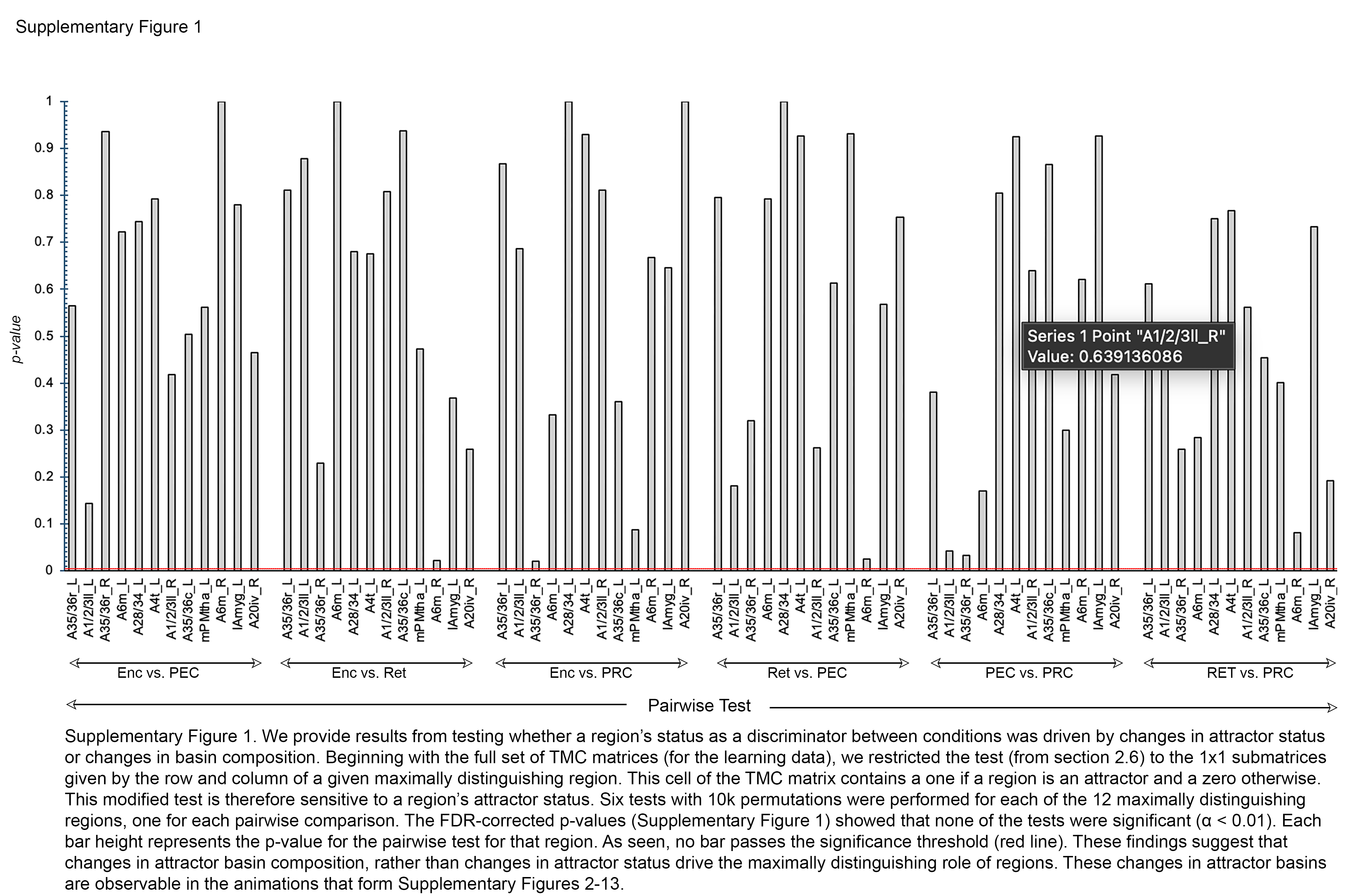
